# Inferred interactions transform values into lattice-patterned choice attractors in orbitofrontal cortex

**DOI:** 10.64898/2026.11.23.768769

**Authors:** Kaining Zhang, Yuma Kanazawa, Timothy G. Crimmins, Camillo Padoa-Schioppa, Gaia Tavoni

## Abstract

Economic decisions require transforming option values into choices, a process involving the orbitofrontal cortex (OFC), but the circuit mechanism remains unclear. We recorded OFC activity in monkeys performing a juice-choice task, fitted graphical Ising models to spike data, and simulated the inferred networks. Offer inputs shifted the networks toward attractor states favoring the choice representation supported by relative value. Targeted ablations revealed a core functional backbone for this transformation, linking offer-value and choice-tuned populations, alongside amplification by broader network interactions and complementary contributions from positive- and negative-encoding pathways. Mean interactions between tuning groups captured the average transformation, while finer interaction structure shaped its network-specific implementation. At single-neuron resolution, the transformation was expressed through a recurring response motif coupling activation and suppression in a spatial lattice. Together, these results reveal how inferred OFC interactions transform offer values into choice-selective attractor states and shape the lattice-patterned neuronal recruitment characterizing this transformation.

## Introduction

Economic decision-making involves transforming representations of option values into a choice. Converging evidence from clinical studies [1–6], lesion analyses [7, 8], neurophysiology [9–12], and functional imaging [13, 14] implicates the orbitofrontal cortex (OFC) in this process. Consistent with this view, OFC microstimulation in both humans [15] and monkeys [16, 17] selectively disrupts value-based decisions. Electrophysiological recordings in primates performing value-based choice tasks further show that OFC activity encodes key decision variables, including the values of offered options and the identity of the chosen good [18–22]. However, how value signals are transformed into reliable choices remains unclear. One prominent hypothesis is that decisions arise from transitions between attractor states in recurrent neural networks [23–28]. In this framework, different choices correspond to distinct attractors in an energy landscape shaped by synaptic interactions and external inputs. Offer values are hypothesized to reshape this landscape: graded changes in relative offer values would progressively alter the accessibility and stability of the competing attractors and drive population activity from a baseline state toward the favored choice-specific attractor. Such a shift involves coordinated changes in the responses of many neurons. We refer to this input-driven, collective reorganization of equilibrium population activity as an attractor transition, or avalanche [29], without implying discontinuous switching (Fig. 1A).

**Figure 1.**
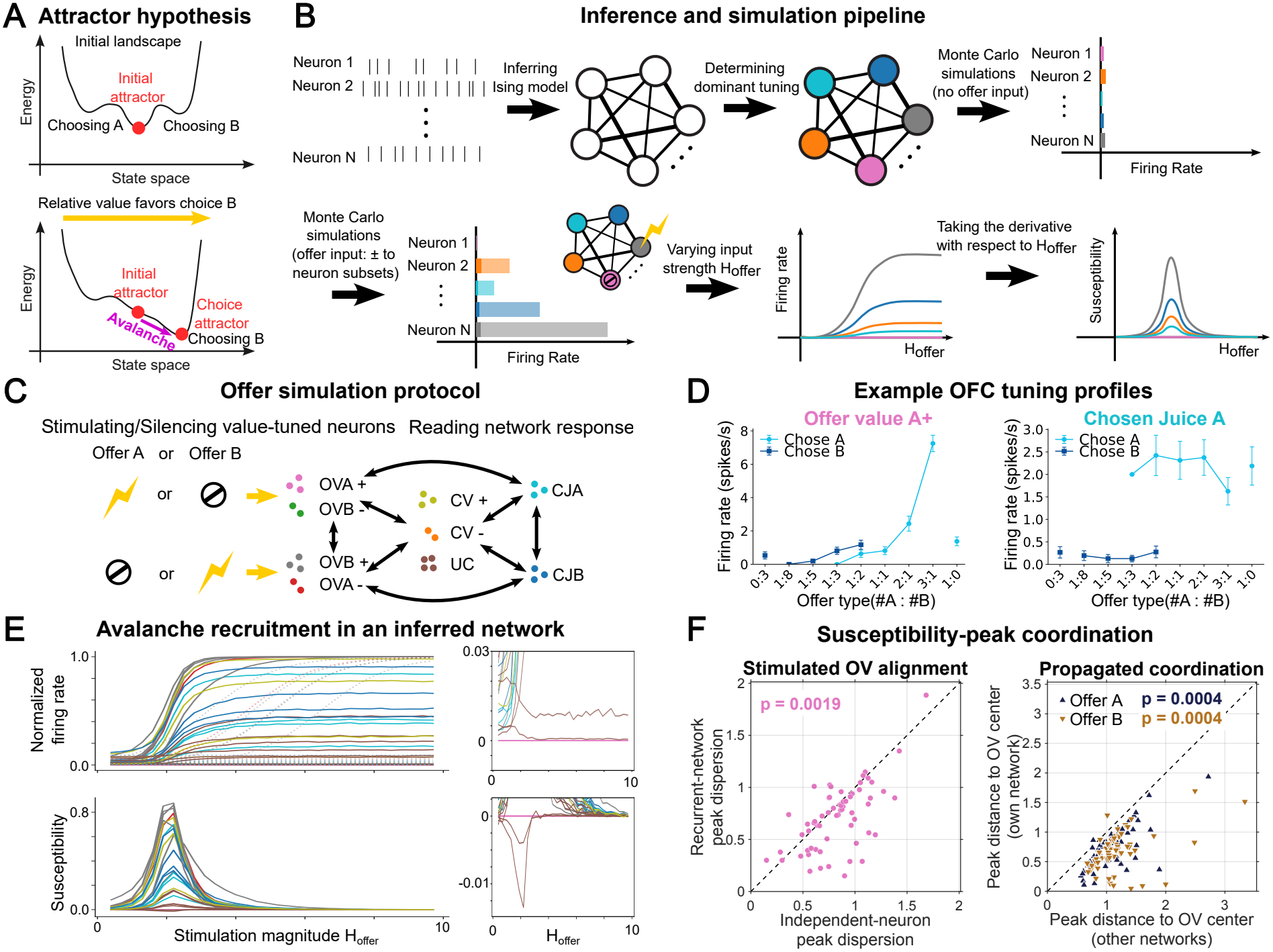
Network inference and simulation framework. (**A**) Attractor hypothesis of decision-making. Network interactions and external inputs shape an energy landscape in which low-energy states represent attractors. Offer presentation reshapes this landscape, driving the network from an initial attractor (top) toward a choice-related attractor (bottom). (**B**) Computational analysis pipeline. We inferred a maximum-entropy Ising model from neural spike data and labeled neurons by the task variable they predominantly encoded (colors; see main text). The inferred networks were simulated using Monte Carlo sampling to characterize equilibrium activity states. A single offer was simulated by applying a positive input *H*_offer_ (lightning-bolt symbol) and a strong negative input (stop sign) to complementary OV populations (e.g., gray and pink), representing the presence of one offer and the absence of the alternative. As *H*_offer_ increases, activity propagates beyond stimulated neurons, changing firing rates in other populations (e.g., blue and orange) relative to baseline (no input). We quantified these responses using neuronal susceptibility, the derivative of firing rate with respect to input strength. Alignment of susceptibility peaks across neurons indicates a collective shift between attractor states, while the Avalanche Index (AI), defined as the susceptibility peak corrected for the direct effect of external input, quantifies the sign and strength of each neuron’s network-mediated response. The schematic shows one neuron per tuning group, but simulations were performed on the full inferred networks at single-neuron resolution. (**C**) Schematic of an offer simulation. For offer A, we stimulated OVA+ and OVB *−*neurons and silenced OVB+ and OVA *−*neurons; for offer B, we reversed these inputs. We then measured the responses of all neurons, including CJA and CJB. (**D**) Firing profiles (mean*±* SEM across trials, separately for choices A and B) of two example neurons recorded in OFC (OVA+ and CJA). Across offer types, the OVA+ neuron exhibits monotonic value coding, with activity increasing with the quantity of juice A, whereas the CJA neuron exhibits approximately binary choice coding, becoming active when juice A is chosen. (**E**) Top: Firing rates (spikes per 5-ms time bin) obtained by Monte Carlo simulations of an example inferred network under offer B, plotted as a function of input strength *H*_offer_. Dotted lines show independent-neuron responses (obtained by setting *J*_*ij*_ = 0). Network interactions tend to coordinate neuronal responses by aligning the input strengths at which firing rates change most steeply. Bottom: Corresponding susceptibilities exhibit aligned positive and negative peaks. Insets to the right show suppressed neurons on an enlarged scale. Colors indicate dominant tuning, as in (C). (**F**) Susceptibility-peak coordination. Left: Susceptibility-weighted dispersion of peak input strengths among stimulated OV neurons is smaller in inferred than in independent-neuron networks (*J*_*ij*_ = 0). Each point represents one network, averaged across offers (*p* = 0.0019, Wilcoxon signed-rank test). Right: In nearly all networks (61*/*62 for each offer), non-stimulated neurons’ peaks are, on average, closer to the stimulated OV peak center of their own network than to those of other networks (both *p*_Holm_ = 4.0 *×*10^*−*4^, permutation tests). Dashed lines indicate equality.

Some models place the decision process within OFC itself [30], whereas others propose that OFC value signals are transformed into choices downstream through accumulation-to-threshold or competitive attractor-like dynamics [26, 31–33]. These frameworks capture important behavioral and neural phenomena but rely on high-level dynamical descriptions or assumed circuit architectures and learning mechanisms, rather than inferring pairwise interaction structure directly from neural data [23, 25, 30, 32, 33]. Population recordings have meanwhile revealed patterns compatible with attractor processes in OFC and PFC, including rapid transitions between option representations, steeper population flow fields for more consistent decisions, low-dimensional trajectories converging onto discrete choice states, and task-specific attractor-like dynamics associated with belief-state updating [21, 22, 34–40]. However, these studies characterize population dynamics and leave open whether and how the effective interaction structure of the recorded populations can itself generate choice-selective attractor states.

Here, we addressed this question by inferring the effective interaction structure of OFC populations directly from spike data and probing its computational implications. We inferred a pairwise maximum-entropy (Ising) model from simultaneously recorded OFC neurons in monkeys performing a value-based choice task, with one model fitted per recording session. Ising models have been used in two parallel traditions in neuroscience: as models of associative memory, where interactions encode attractor states in the Hopfield formulation [41–43]; and as least-biased descriptions of recorded population activity consistent with measured firing rates and pairwise correlations, used to investigate collective properties. The latter tradition has demonstrated that when fitted to spiking data, the Ising model exhibits powerful generative properties, accurately predicting population activity patterns across a variety of brain regions [44–53], including prefrontal and orbitofrontal cortex [29, 54, 55].

In this study, we combine these two approaches by inferring the model from OFC spike data and using it to investigate the resulting attractor landscape. Specifically, the model provides a principled way to test whether offer inputs acting through the inferred pairwise interactions shift equilibrium population activity from its baseline configuration toward choice-consistent attractor states, and to characterize the internal organization of these transitions. We simulate equilibrium network responses to offer inputs by Monte Carlo sampling of the inferred Ising distributions [56, 57]. We quantify each neuron’s response by its susceptibility, defined as the derivative of its spiking probability with respect to the input. Positive and negative susceptibility extrema mark the input levels at which each neuron’s activity increases or decreases most rapidly, respectively. Their alignment across neurons identifies avalanches – coordinated activation and suppression associated with attractor transitions [29].

We find that offer inputs drive attractor transitions that preferentially activate choice neurons corresponding to the simulated offer and, under competing offers, on average favor those corresponding to the higher-value option. Compared with matched random networks, the inferred networks generate stronger, more coordinated, and more choice-selective avalanches. Targeted ablations show that removing interactions between offer-value and choice-tuned populations nearly abolishes the choice bias, identifying the core of the transformation. Broader network interactions amplify the effect, while positive- and negative-encoding pathways make complementary, largely additive contributions. Tuning-based coarse-graining of the interaction structure preserves the direction and much of the magnitude of the value-to-choice mapping on average across networks, but poorly reproduces individual network choice biases. Thus, the transformation can be understood at two scales: mean functional interactions capture its aggregate behavior, whereas finer interaction structure shapes its network-specific expression and neuron-resolved implementation. At this finer scale, the transformation involves more than the uniform activation or suppression of tuning-defined populations. Neurons with similar tuning instead split into activated and suppressed subpopulations. These opposing recruitment roles are not arranged arbitrarily in space: local co-activation is coupled to offset suppression in a characteristic motif that recurs statistically across recording sites, defining a spatial lattice of neuronal recruitment.

Together, these findings identify the roles of different interactions in shaping value-to-choice attractor transitions in OFC and reveal how the resulting offer-dependent neuronal recruitment is organized within tuning groups and across space.

## Results

### Behavioral task and neuronal recordings

Three male monkeys performed a binary economic choice task described previously [55] and detailed in the Methods. In each session, they were offered varying quantities of two juices, A and B, with A preferred, represented as colored squares on a monitor. After a delay, the monkeys indicated their choice with a saccade, following which the chosen juice was delivered. The monkeys’ decisions reflected a trade-off between juice preference and quantity, and the session-specific relative value of the two juices, *ρ*, was estimated from their choice patterns.

Extracellular recordings targeted OFC area 13m and were performed across 85 sessions using 32-channel linear probes with contacts spaced by 75 *µ*m. One to four probes were used per session, with at most two probes per hemisphere, allowing simultaneous sampling of neuronal populations across cortical depth and recording sites. Of the 85 sessions, 63 met predefined inclusion criteria for the computational analyses (Methods). Each included session contained 19–116 simultaneously recorded neurons (median = 53).

### Network inference and simulation framework

We analyzed this dataset using the computational pipeline illustrated in Fig. 1B. For each session, we fitted an Ising model to the firing rates and pairwise correlations of simultaneously recorded neurons (Methods). The Ising model is the maximum-entropy, or least-constrained, model consistent with these statistics. Its parameters describe the pairwise effective interactions between neurons, *J*_*ij*_, and each neuron’s intrinsic bias toward activity, *h*_*i*_. The model defines a probability distribution over population activity states, (*σ*_1_, … , *σ*_*N*_), where *σ*_*i*_ = 1 if neuron *i* is active in a given 5-ms time bin and *σ*_*i*_ = 0 otherwise. Unless otherwise specified, one model was inferred per session using all trials, spanning different offer quantities and choices.

We then probed the equilibrium responses of the inferred networks to inputs representing different offers. In the absence of external drive, the inferred networks generally occupy low-activity states. We therefore applied a uniform baseline input to all neurons to place each network in an operating regime from which selective perturbations could reveal whether the inferred interaction structure supports collective transitions. To simulate different offers, we then applied selective inputs to neurons predominantly tuned to the corresponding offer values. Specifically, we identified neurons with the following dominant tuning properties [20, 55]: OVA+ and OVA*−* neurons increase or decrease their firing rates with offer value A, respectively; OVB+ and OVB*−* neurons are defined analogously for offer value B; CV+ and CV*−* neurons covary positively or negatively with the chosen value, respectively; and CJA and CJB neurons increase their firing rates when juice A or B is chosen. Neurons for which a dominant tuning could not be identified were labeled UC (unclassified). Fig. 1D shows the firing profiles of two example neurons. To simulate the presentation of offer A alone, we applied a positive input, *H*_offer_ = *H*_*A*_, to OVA+ and OVB*−* neurons, corresponding to the presence of A and the absence of B. Correspondingly, we applied a strong negative input to OVB+ and OVA*−* neurons, suppressing populations that signal the presence of B and the absence of A. Similarly, to simulate the presentation of offer B alone, we applied a positive input, *H*_offer_ = *H*_*B*_, to OVB+ and OVA*−* neurons and a strong negative input to OVA+ and OVB*−* neurons (Fig. 1C). We also simulated intermediate conditions by applying both *H*_*A*_ and *H*_*B*_ simultaneously to their respective populations, with a combined offer input amplitude *H*_total_ = *H*_*A*_ + *H*_*B*_. Ratios *H*_*B*_*/H*_*A*_ *>* 1 favored offer B, whereas ratios *H*_*B*_*/H*_*A*_ *<* 1 favored offer A. Importantly, inputs *H*_*A*_ and *H*_*B*_ were not intended to reproduce the full stimulus-evoked activity pattern associated with each offer. Instead, we used first-order, offer-aligned perturbations to test whether the inferred network could amplify the value input and transform it into a choice-consistent state. In this spirit, restricting stimulation to neurons whose dominant tuning is aligned with a given offer provides a controlled way to probe how activity propagates through the inferred interaction structure. Neurons with secondary tuning to that offer were not directly stimulated but could be recruited through network interactions.

In each simulation, we gradually varied the offer input strength (*H*_*A*_, *H*_*B*_, or *H*_total_, depending on the condition). The firing rates of the directly stimulated OV neurons increased nonlinearly, accompanied by increased activity in other neurons and decreased activity in a smaller subset (Fig. 1E, top). To systematically characterize how this offer perturbation propagated through the network, we computed each neuron’s susceptibility, defined as the derivative of its equilibrium firing rate with respect to input strength (Fig. 1E, bottom). Positive and negative susceptibility peaks quantify the maximal rates of increase and decrease in neuronal activity, respectively, as input strength varies. Input-driven shifts between attractor states, or avalanches, are characterized by coordinated, network-mediated neuronal responses, identified by susceptibility peaks occurring at similar input strengths [29].

To assess this coordination across inferred networks, we first compared the dispersion of input strengths at which susceptibility peaks occur among stimulated OV neurons with the corresponding independent-neuron predictions (*J*_*ij*_ = 0), in which neuronal responses depend only on intrinsic biases and external inputs. Peak dispersion is significantly smaller in the inferred networks (median paired difference = *−*0.055, 95% CI [*−*0.121, *−* 0.018]; *p* = 0.0019, Wilcoxon signed-rank test; Fig. 1F, left), indicating that recurrent interactions promote the alignment of stimulated neuronal responses. We next examined whether this coordination extends to non-stimulated neurons. In nearly all networks (61*/*62 for each offer), susceptibility peaks are, on average, closer to the stimulated OV peak center of their own network than to those of other networks. Across sessions, the mean distance is 33.2% smaller for offer A and 43.0% smaller for offer B compared with mismatched network centers (median coordination effect *E* = *−*0.273, 95% CI [*−*0.333, *−*0.231] for A; *E* = *−*0.363, 95% CI [*−*0.465, *−*0.312] for B; both *p*_Holm_ = 4.0*×*10^*−*4^, permutation tests; Fig. 1F, right; Methods). Together, these results demonstrate systematic network-specific coordination of susceptibility peaks across directly stimulated and propagated responses.

Having established this population-level coordination, we sought to understand how the inferred interactions within OFC networks shape individual neuronal recruitment during attractor transitions. Because direct stimulation can generate susceptibility peaks even without network interactions, we defined the Avalanche Index (AI) as the signed value of each neuron’s susceptibility peak (maximum or minimum) after subtracting the maximal direct-input contribution expected for independent neurons (Methods). AI thus quantifies network-mediated changes in neuronal responsiveness, allowing us to determine which neurons participate in avalanches and how strongly.

### Offer inputs trigger attractor transitions toward offer-aligned choice states in OFC networks

For each inferred network, we separately simulated offers A and B. Because nonzero AI can arise even with randomized *J*_*ij*_, and AI magnitude also depends on global network properties (Fig. 2D), we compared each inferred network with random controls matched for network size, number of stimulated neurons, and the means and variances of *J*_*ij*_ and *h*_*i*_, but with otherwise random parameters (Methods).

**Figure 2.**
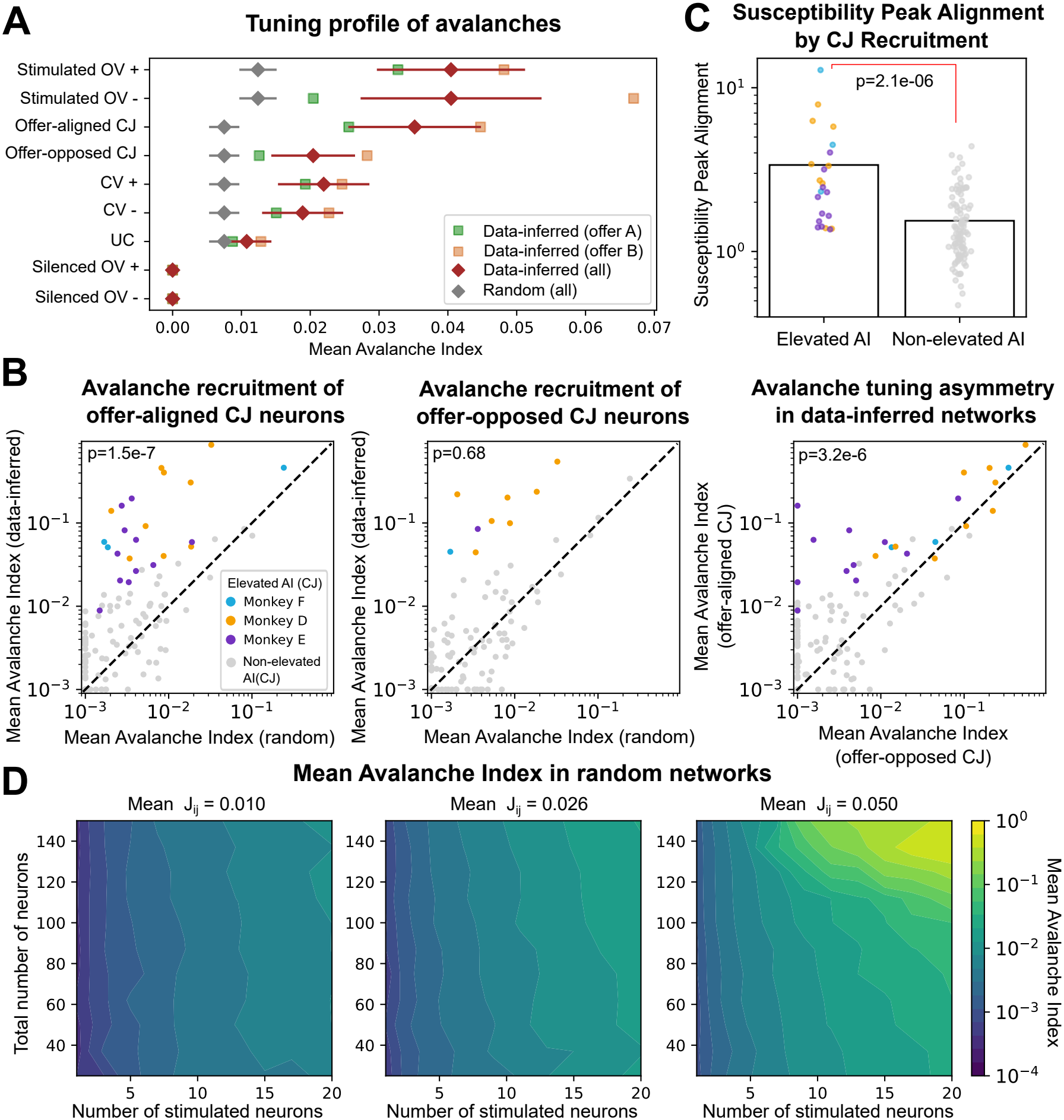
Offer inputs trigger attractor transitions toward offer-aligned choice states in OFC networks. (**A**) Tuning profile of avalanches. In data-inferred networks, neurons are grouped by dominant tuning, and AI is averaged within each group. Green and yellow data points show session-averaged values for offer A and offer B simulations, respectively; red data points and bars show the mean*±* SEM across all sessions and simulation conditions. In matched random networks (gray), values are computed separately for stimulated and non-stimulated neurons (Methods). All cell groups show higher AI in data-inferred than in matched random networks, with offer-aligned CJ neurons showing the highest AI among non-stimulated groups. (**B**) Avalanche recruitment of CJ neurons. Left and middle: Mean AI of offer-aligned and offer-opposed CJ neurons, respectively, in data-inferred networks versus mean AI of non-stimulated neurons in matched random networks. Right: Mean AI of offer-aligned versus offer-opposed CJ neurons in data-inferred networks. Each point represents one session under one offer simulation. Blue, orange, and purple indicate, for each monkey, conditions with elevated mean AI of offer-aligned CJ neurons (left), offer-opposed CJ neurons (middle), or either type (right); gray indicates non-elevated conditions; *p*-values are from Wilcoxon signed-rank tests comparing x- and y-axis values. Values are clipped at 10^*−*3^ for visualization only. Avalanches predominantly recruit offer-aligned CJ neurons. (**C**) Susceptibility Peak Alignment in conditions with elevated versus non-elevated mean AI of CJ neurons. Each point represents one session under one offer simulation; colors are as in (B), right. Alignment is higher in elevated than non-elevated conditions (*p* = 2.1*×* 10^*−*6^, Wilcoxon rank-sum test), indicating tighter population-level coordination when CJ recruitment is stronger. (**D**) Mean AI of non-stimulated neurons in randomly connected networks as a function of network size, number of stimulated neurons, and mean interaction strength, *⟨J*_*ij*_*⟩*. The standard deviation of *J*_*ij*_ is fixed at the dataset-wide value of 0.22; the central plot uses the dataset-wide mean of *J*_*ij*_. For each parameter combination, 50 random realizations were generated and results were averaged across those satisfying the Monte Carlo convergence criterion (Supplementary Information). Mean AI increases with all three parameters, although at different rates.

We first asked whether the inferred interactions promote greater susceptibility-peak coordination than matched random networks. Susceptibility Peak Alignment is significantly higher in inferred networks (*p* = 0.0054, Wilcoxon signed-rank test; Fig. S1), supporting a role for the inferred interaction structure in coordinating population responses.

We then compared mean AI across tuning groups between inferred and matched random networks, averaging across sessions and both offer simulations (Fig. 2A). Inferred-network values are also shown separately for each offer. We define offer-aligned CJ neurons as CJA during offer A simulations and CJB during offer B simulations; the alternative CJ population is termed offer-opposed.

Fig. 2A reveals three main findings. First, mean AI is generally higher in data-inferred than in matched random networks (with a significant difference, *p <* 0.05, Wilcoxon signed-rank test, in all groups except offer-opposed CJ and OV*−* neurons), indicating that the specific structure of OFC interactions promotes offer-evoked avalanche recruitment. Second, offer-aligned CJ neurons exhibit the highest mean AI among non-stimulated groups, indicating preferential recruitment. Third, these patterns remain qualitatively unchanged when mean AI is analyzed separately for offer A and offer B simulations.

We next examined the mean AI of CJ neurons in individual conditions, each corresponding to one session under one offer simulation (Fig. 2B). Some conditions show elevated mean AI, defined as more than 3 SEM above the matched-random mean, for either offer-aligned or offer-opposed CJ neurons. However, every condition with elevated offer-opposed CJ AI also shows comparably high offer-aligned CJ AI, whereas the converse is not true. Moreover, mean AI is significantly higher for offer-aligned than for offer-opposed CJ neurons across conditions (*p* = 3.2*×* 10^*−*6^, Wilcoxon signed-rank test). Thus, avalanches typically exhibit a tuning asymmetry favoring recruitment of offer-aligned CJ neurons.

Finally, conditions with stronger recruitment (elevated mean AI) of CJ neurons also show tighter peak alignment (*p* = 2.1 *×*10^*−*6^, Wilcoxon rank-sum test), linking stronger value-to-choice recruitment to a more coordinated population response (Fig. 2C).

In conclusion, the inferred OFC interaction structure propagates offer inputs into coordinated avalanche responses with a net activation bias and preferential recruitment of choice neurons aligned with the simulated offer. These responses reflect input-driven shifts toward the corresponding choice-selective attractor states.

### Competing offers bias attractor transitions toward the higher-value choice

In the previous section, we examined OFC network responses to single offers. Here, we examine responses when two offers are presented simultaneously. For simplicity, we applied the same input drive *H*_*A*_ to OVA+ and OVB*−* neurons, and the same input drive *H*_*B*_ to OVB+ and OVA*−* neurons, simulating conditions in which the two offers occupied approximately complementary positions within their respective value ranges (Methods). The ratio *H*_*B*_*/H*_*A*_ (input bias) was held fixed within each simulation and varied across simulations. Ratios *H*_*B*_*/H*_*A*_ *>* 1 simulated offer conditions in which B had higher value than A, whereas ratios *H*_*B*_*/H*_*A*_ *<* 1 simulated conditions in which A had higher value than B. For each session and input bias, we defined Avalanche Asymmetry as the mean AI of CJB neurons minus the mean AI of CJA neurons. Avalanche Asymmetry thus quantifies the choice bias of the value-to-choice transformation, measuring the preferential recruitment of neurons representing choice B versus choice A.

Fig. 3A shows the mean Avalanche Asymmetry across sessions. Notably, when *H*_*B*_*/H*_*A*_ = 1 (symmetric input, corresponding to equal offer values for A and B), the mean Avalanche Asymmetry is statistically indistinguishable from zero (*p* = 0.15, Wilcoxon signed-rank test). As the input bias *H*_*B*_*/H*_*A*_ increases above 1, the mean Avalanche Asymmetry increases, indicating that, on average, the networks transition toward attractor states that more strongly recruit CJB than CJA neurons. Conversely, as the input bias *H*_*B*_*/H*_*A*_ decreases below 1, the mean Avalanche Asymmetry becomes negative, reflecting preferential recruitment of CJA neurons. Thus, the average network response is systematically biased toward the choice representation associated with the higher-value offer.

**Figure 3.**
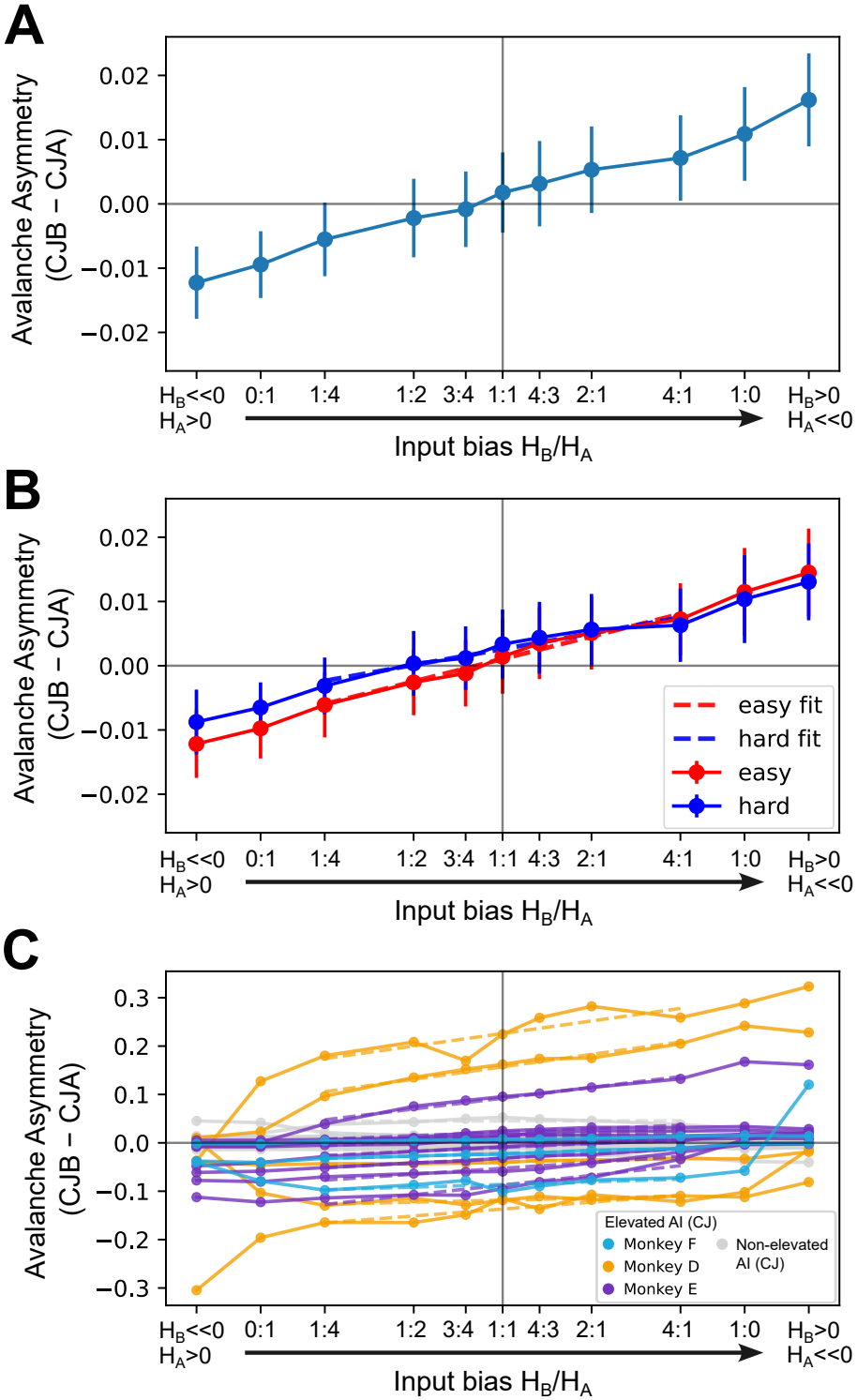
Competing offers bias attractor transitions toward the higher-value choice. (**A**) Avalanche Asymmetry in the fixed CJB *−* CJA co-ordinate (AA_CJB*−*CJA_), mean *±* SEM across sessions, as a function of the input bias *H*_*B*_*/H*_*A*_ (x-axis in log-scale except for the two leftmost and two rightmost values). The leftmost and rightmost points correspond to the previous offer-A-only and offer-B-only simulations. As the input bias shifts from favoring offer A (*H*_*B*_*/H*_*A*_ *<* 1) to favoring offer B (*H*_*B*_*/H*_*A*_ *>* 1), mean Avalanche Asymmetry changes from negative to positive, indicating a corresponding shift from CJA-dominated to CJB-dominated avalanche recruitment. (**B**) Same as in (A), but for networks inferred separately from easy (red) and hard (blue) trials. Mean Avalanche Asymmetry increases with log(*H*_*B*_*/H*_*A*_) in both sets of networks, with no significant difference between the fitted slopes (*β*_easy_ = 0.0117, *β*_hard_ = 0.0082; *p* = 0.13, Wald test). (**C**) Same analysis as in (A), shown for individual sessions. Colored curves indicate, for each monkey, sessions with elevated mean AI of CJA or CJB neurons in the two extreme offer conditions, as in Fig. 2; gray curves indicate the remaining sessions. Individual networks show different intrinsic biases toward choice A or B, reflected by non-zero Avalanche Asymmetry at *H*_*B*_*/H*_*A*_ = 1. Despite this variability, Avalanche Asymmetry generally increases with *H*_*B*_*/H*_*A*_, indicating shifts toward attractor states favoring choice B as its relative offer input increases.

We next examined whether this result depended on the behavioral choice difficulty of the trials used to infer the networks. We split the recorded trials into behaviorally easy and hard decisions based on the relative value difference between the juices offered to the monkey (Methods), re-inferred the Ising model separately from each subset, and simulated the resulting networks using the same protocol. In both cases, mean Avalanche Asymmetry increased similarly with input bias (Fig. 3B), with no significant difference between the fitted slopes (*p* = 0.13, Wald test).

Finally, we analyzed Avalanche Asymmetry at the level of individual sessions. Fig. 3C reveals substantial session-to-session variability. In particular, some networks exhibit non-zero Avalanche Asymmetry at *H*_*B*_*/H*_*A*_ = 1, indicating an intrinsic bias toward choice A or B even under symmetric stimulation. This variability is expected because each session samples only a limited subset of the underlying OFC population, capturing a sparse random projection of the larger biological circuit. Importantly, even biased networks typically show increasing Avalanche Asymmetry with increasing input bias. For each session, we fitted Avalanche Asymmetry as a linear function of log(*H*_*B*_*/H*_*A*_) across the non-extreme competing-offer conditions, from *H*_*B*_*/H*_*A*_ = 1 : 4 to 4 : 1. 48*/*61 sessions (78.69%) have positive slopes, with 34*/*61 (55.74%) reaching significance (*p <* 0.05, Wald test), compared to only 6*/*61 (9.84%) with a significantly negative slope.

Together, these results demonstrate that inferred OFC networks transform relative differences in offer value into shifts toward the choice attractor associated with the higher-value option, while remaining unbiased on average for symmetric offers.

### Network ablations and coarse-graining reveal the functional architecture of the value-to-choice transformation

To identify which components of the inferred interaction structure are responsible for transforming offer inputs into recruitment of the corresponding choice populations, we performed two complementary network-reduction tests: neuron-resolved ablations and coarse-grained reductions. The ablations ask which specific pathways are necessary for, or contribute to, the transformation by selectively removing targeted interactions while leaving the rest of the neuron-resolved network unchanged. The coarse-grained reductions instead ask whether the detailed neuron-to-neuron interaction structure can be compressed into a lower-dimensional description while preserving the equilibrium value-to-choice transformation.

#### Core OV–CJ interactions drive the value-to-choice transformation, with complementary encoding pathways and amplification by the broader network

For each ablated network, we repeated the offer A/B simulations of Fig. 2 and quantified offer-aligned Avalanche Asymmetry as the difference between the mean AI of offer-aligned and offer-opposed CJ neurons (Fig. 4). Fig. 4A shows the effects of ablating specific functional groups or interactions between them. Removing only OV–CJ interactions (“OV/CJ–decoupled network”) dramatically reduces Avalanche Asymmetry, despite eliminating only a small fraction of couplings. The impact is substantially stronger than that of random ablations removing the same number of couplings (*p* = 3.3 *×*10^*−*7^, Wilcoxon rank-sum test). By contrast, removing only UC neurons from the full inferred networks (“canonical task network”) causes a modest reduction in Avalanche Asymmetry, despite eliminating a large fraction of the network. The “OV–CJ core network”, obtained by additionally removing CV neurons, still outperforms matched random controls (*p* = 1.3 *×*10^*−*7^, Wilcoxon rank-sum test) and retains *∼*60% of the full-network Avalanche Asymmetry. These results indicate that the OV–CJ circuit is sufficient to support a substantial fraction of the transformation and forms its core functional architecture, whereas CV and UC populations primarily amplify its magnitude. Fig. 4B examines the positive-encoding pathway (OVA+, OVB+, CV+) and negative-encoding pathway (OVA*−* , OVB*−* , CV*−*), each retained separately with CJ neurons or combined without cross-pathway interactions (“parallel-encoding network”). Both pathways independently generate Avalanche Asymmetry, with a stronger contribution from the positive pathway. Combining them without cross-pathway interactions produces an approximately additive effect, while restoring such interactions provides only modest additional amplification. Thus, the two pathways offer complementary, largely independent routes for the value-to-choice transformation, with modest synergistic reinforcement from their mutual interactions.

**Figure 4.**
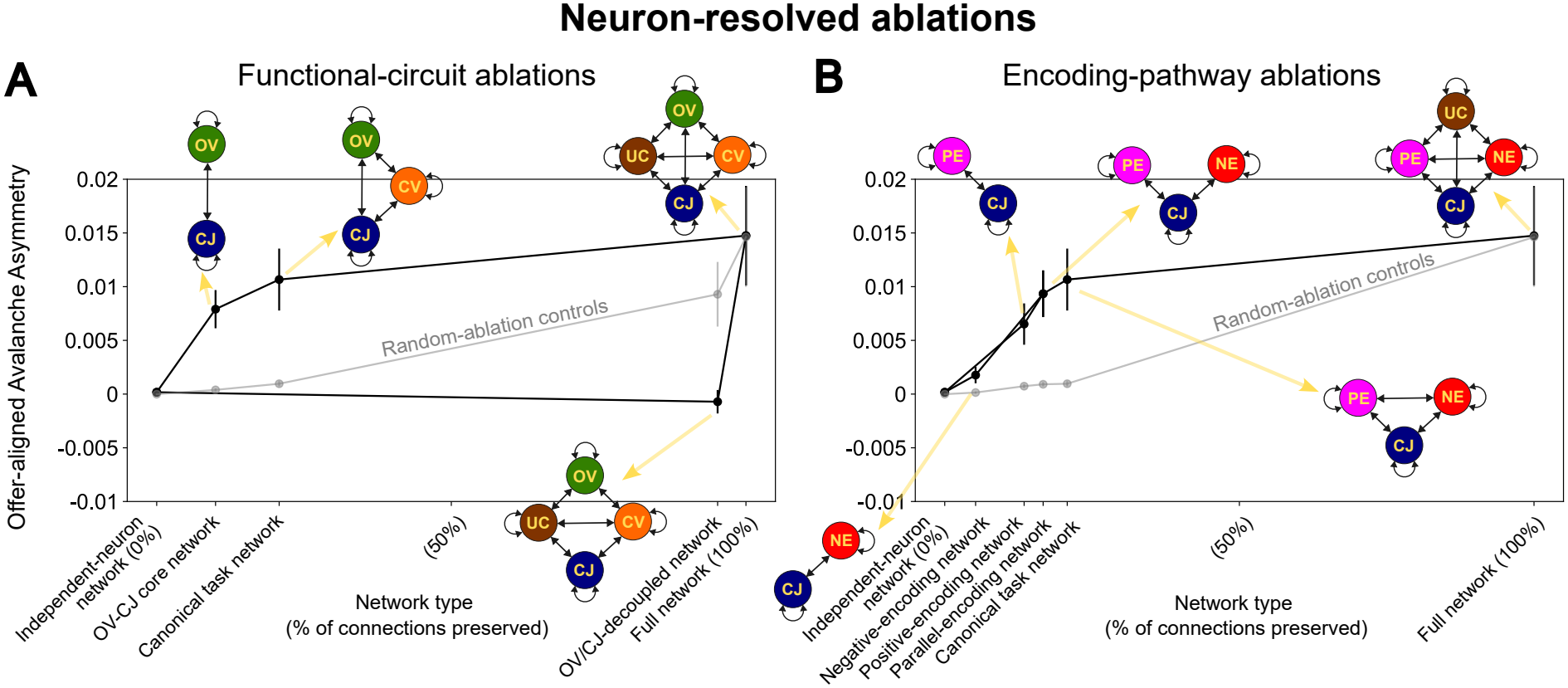
Core OV–CJ interactions drive the value-to-choice transformation, with complementary encoding pathways and amplification by the broader network. Selected neuronal populations or interaction classes were removed directly from the full inferred networks, while all remaining pairwise interactions *J*_*ij*_ and neuronal biases *h*_*i*_ were left unchanged. For each ablated network, the same offer simulations as in Fig. 2 were repeated. The x-axis indicates the fraction of the full network preserved and the corresponding subnetwork type. Black points show mean *±*SEM offer-aligned Avalanche Asymmetry across sessions and offer simulations; gray points show matched random ablations removing the same number of connections uniformly across neuron pairs. (**A**) Removing OV–CJ interactions nearly abolishes Avalanche Asymmetry, whereas the OV–CJ core alone retains *∼*60% of its full-network magnitude. The CV and UC populations provide further amplification. (**B**) Positive-encoding (PE) and negative-encoding (NE) pathways, together with the CJ population, each generate Avalanche Asymmetry, with a stronger contribution from the positive pathway. Combining the two pathways without cross-pathway interactions produces an approximately additive effect, indicating largely independent contributions, whereas restoring cross-pathway interactions provides only modest additional amplification.

#### Coarse functional architecture captures the aggregate value-to-choice transformation, while neuron-resolved interactions shape its network-specific realization

For each inferred network and single- or competing-offer condition, we next computed Avalanche Asymmetry in two successive coarse-grained models: a functional-group mean-field (FG-MF) reduction that replaces neuron-to-neuron couplings by group-level means, and a two-channel mean-field (2C-MF) reduction that further retains only the mean group-to-group couplings along the offer-aligned OV–CJ channels, i.e., those linking OVA+ and OVB*−* to CJA, and OVB+ and OVA*−* to CJB. For each reduction, we considered a neuron-biased version retaining the inferred *h*_*i*_ and a group-biased version replacing them by tuning-group means *h*_*u*_ (Methods).

We first studied the transformation at the aggregate level, i.e., its average functional expression across networks. In all reduced models, mean Avalanche Asymmetry varies systematically with the *H*_*B*_*/H*_*A*_ input bias, indicating preservation of the choice-aligned direction of the transformation and part of its magnitude. Its magnitude is progressively attenuated as the interaction structure is compressed, while further compression of neuron-specific biases *h*_*i*_ to group means *h*_*u*_ has little effect. Across the 10 input conditions with a favored option, the neuron- and group-biased FG-MF reductions preserve 67% and 68% of the full-network asymmetry, respectively, while the corresponding 2C-MF reductions preserve 51% and 53% (Fig. 5A). Thus, all four reductions preserve much of the aggregate mapping from relative offer input to choice-population recruitment.

**Figure 5.**
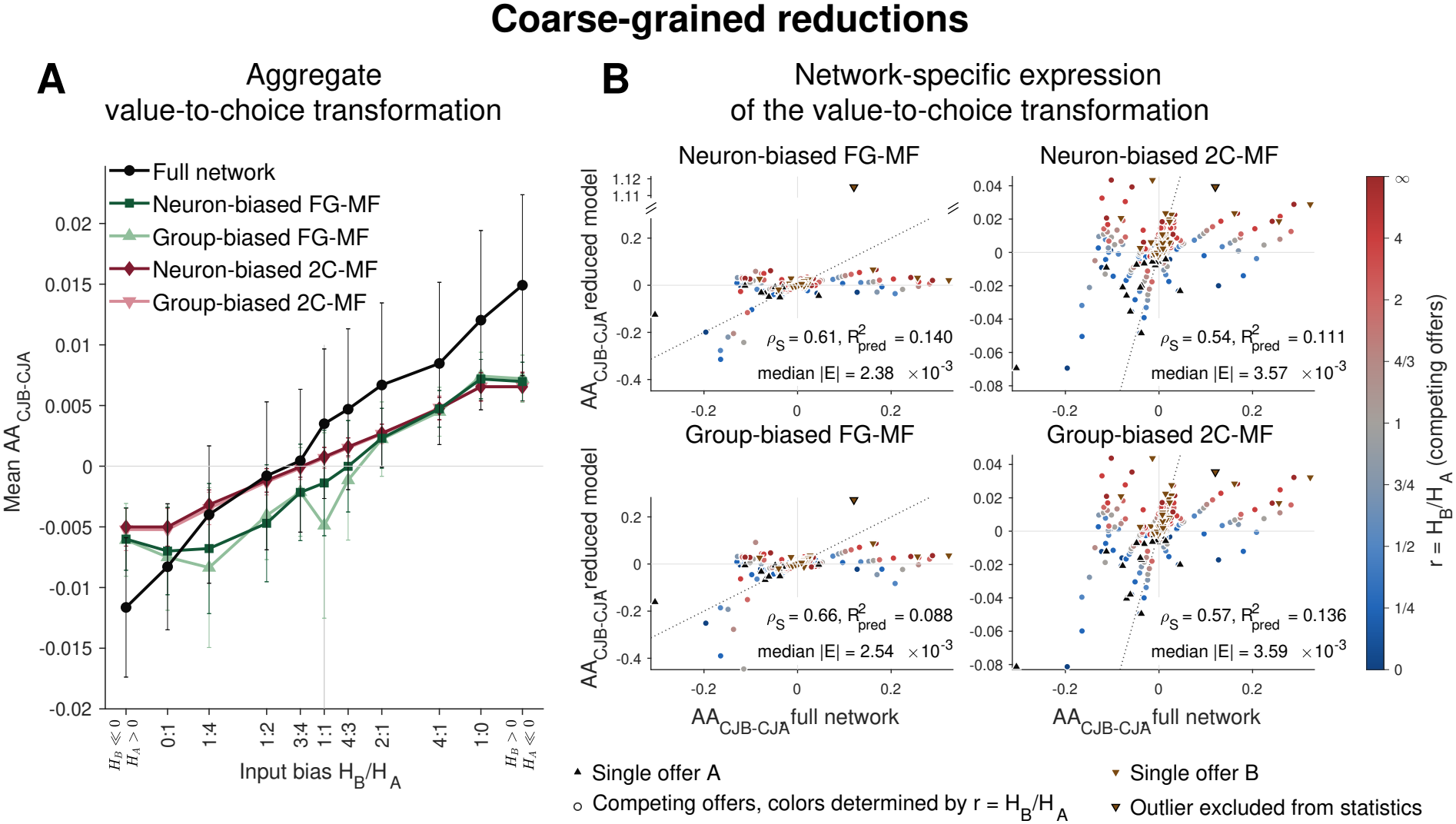
Coarse functional architecture captures the aggregate value-to-choice transformation, while neuron-resolved interactions shape its network-specific realization. (**A**) Mean *±* SEM Avalanche Asymmetry in the fixed CJB *−* CJA coordinate (AA_CJB*−*CJA_) across sessions is plotted against input bias *H*_*B*_*/H*_*A*_, as in Fig. 3A, for the full network and four coarse-grained reductions (model definitions in the main text). Compared with Fig. 3A, one session is excluded here because it produces an extreme near-singular amplification in the neuron-biased FG-MF (see B). All reductions preserve much of the aggregate functional mapping from relative offer input to choice-recruitment bias when averaged across networks, with generally reduced magnitude relative to the full network. Neuron- and group-biased models perform similarly. (**B**) Network-specific realization of the value-to-choice transformation. Each point compares reduced- and full-network Avalanche Asymmetry in the fixed CJB*−* CJA coordinate for one session–offer condition; dotted lines indicate identity. Triangles denote single offers and circles competing offers, colored by *H*_*B*_*/H*_*A*_. Spearman *ρ*_*S*_ quantifies rank correspondence, 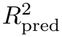 quantifies how well the reduced model predicts the full-network Avalanche Asymmetry relative to simply predicting its mean for every session–offer condition, and median |*E*| quantifies the typical raw prediction error in Avalanche Asymmetry units. All reductions preserve substantial rank ordering but show low 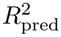, indicating poor recovery of network- and condition-specific response magnitudes. One condition (an outlier in the neuron-biased FG-MF) is excluded from the displayed statistics across all models. Including it leaves rank correspondence essentially unchanged but substantially worsens 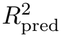 for the neuron-biased FG-MF (from 0.140 to *−*0.433). Together, (A–B) identify a coarse functional backbone that preserves the aggregate mapping, while finer interaction structure is required for its network-specific realization.

However, this aggregate correspondence does not translate into accurate predictions of the transformation within individual networks (Fig. 5B). Consistent with Fig. 5A, across session–offer conditions, all reductions retain substantial rank correspondence with the full-network Avalanche Asymmetry (*ρ*_*S*_ = 0.61–0.66 for the FG-MF models and *ρ*_*S*_ = 0.54–0.57 for the 2C-MF models), but show low 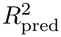 (0.140 for neuron-biased FG-MF, 0.088 for group-biased FG-MF, 0.111 for neuron-biased 2C-MF, and 0.136 for group-biased 2C-MF), indicating poor recovery of the specific choice bias expressed by each network under each input condition. Retaining neuron-specific *h*_*i*_ does not substantially improve this correspondence, indicating that neuronal bias heterogeneity alone cannot account for the missing network specificity. Instead, the network- and condition-specific expression of the transformation depends on neuron-resolved network structure lost when *J*_*ij*_ is replaced by functional-group averages.

Together, these results distinguish two levels of the value-to-choice transformation: a coarse functional level and a neuron-resolved implementation level. Coarse functional interactions, and in particular the offer-aligned OV–CJ backbone, preserve much of the aggregate mapping from relative value to choice recruitment. Finer interaction structure is required to capture the network-specific realization of that mapping. We therefore asked how this finer structure is reflected in the recruitment of individual neurons, and whether it follows a systematic organizational principle.

### Avalanche responses couple activation and suppression within and across functional populations

Here, we examine recruitment at the neuron level. Alongside predominant activation, avalanches also include neurons with negative AI. For directly stimulated OV neurons, negative AI reflects gain suppression – peak responsiveness to the external input below that expected without network interactions, despite increasing activity – whereas for non-stimulated neurons, it reflects activity suppression, with firing decreasing as stimulation increases. Activity-suppressed neurons exhibit inverted bell-shaped susceptibility profiles, with a negative minimum where firing decreases most steeply (Fig. 1E).

This signed heterogeneity is present within tuning groups. Consistent with the positive mean AI in Fig. 2, activated neurons are more prevalent than suppressed neurons (Fig. 6A, left), but both roles coexist: for every tuning group, the mean fraction of neuron pairs with opposite recruitment signs within the same offer simulation ranges from 22.7% to 45.3% (Fig. 6A, right). Thus, dominant functional identity does not uniquely determine a neuron’s role within an avalanche.

**Figure 6.**
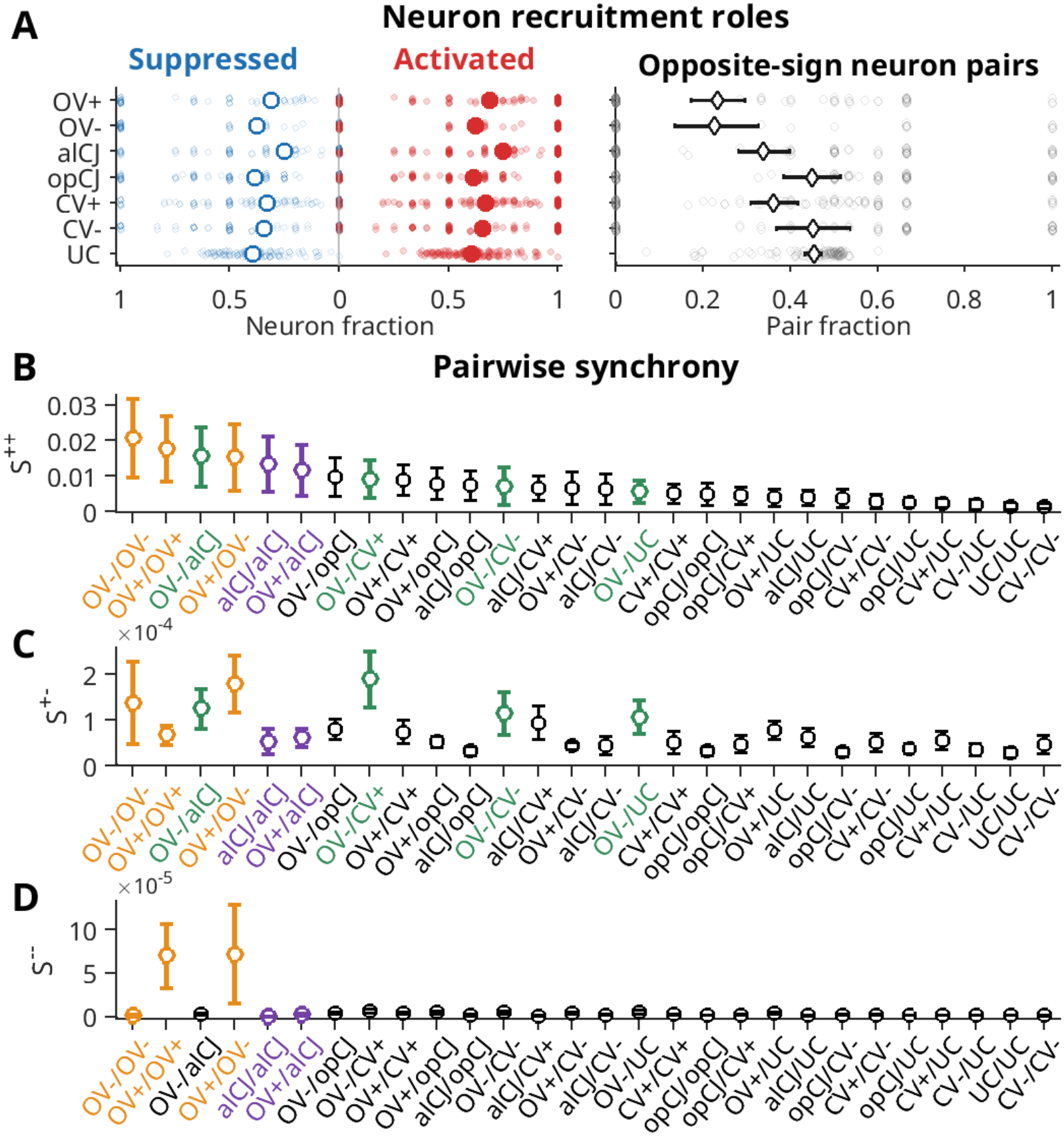
Avalanche responses couple activation and suppression within and across functional populations. (**A**) Neuron recruitment roles within dominant-tuning groups; externally silenced OV groups are excluded from all panels, so OV+ and OV*−* refer to the stimulated groups. Left, fractions of activated (positive AI; filled red circles) and suppressed (negative AI; open blue circles) neurons in each session–offer condition; large symbols show the mean across conditions. Right, fraction of neuron pairs with opposite recruitment signs within the same tuning group and session–offer condition. Diamonds show means across conditions and error bars indicate 95% session-cluster bootstrap confidence intervals. (**B– D**) Synchrony Indices for each pair of dominant-tuning groups. For each tuning-group pair, Synchrony Indices are averaged across constituent neuron pairs within each session–offer condition; points show the mean *±* SEM across conditions. (**B**) *S*^++^ quantifies co-activation, (**C**) *S*^+*−*^ quantifies opposite-sign co-recruitment, and (**D**) *S*^*−−*^ quantifies co-suppression (Methods). Stimulated OV pairs (orange) show prominent same-sign and opposite-sign recruitment. Cross-group pairs involving stimulated OV*−* neurons (green) show elevated opposite-sign recruitment together with intermediate-to-high co-activation. The OV+/offer-aligned CJ and offer-aligned CJ/offer-aligned CJ pairs (purple) combine strong co-activation with little opposite-sign recruitment or co-suppression. Thus, although OV–CJ interactions form the core functional backbone, these patterns show that preferential choice recruitment involves more than a simple activation relay. Rather, stimulation of OV neurons engages a distributed pattern of coordinated activation and suppression, within which the corresponding choice population is preferentially recruited.

We next examined how recruitment roles are organized pairwise within avalanches. We defined three pairwise Synchrony Indices, *S*^++^, *S*^*−−*^, and *S*^+*−*^, quantifying co-activation, co-suppression, and opposite-sign recruitment, respectively (Fig. 6B–D). For each neuron pair, the corresponding index is the product of the two neurons’ AI magnitudes when their signs match the recruitment mode, and zero otherwise (Methods). These indices extend the analysis of Fig. 6A by quantifying the strength of pairwise recruitment within and across tuning groups.

On average, the stimulated OV*− /*OV*−* , OV+*/*OV+, and OV+*/*OV*−* group pairs (orange in Fig. 6B–D) show elevated *S*^++^ together with elevated *S*^+*−*^ or *S*^*−−*^. This signed structure indicates that network interactions enhance input-driven recruitment in some stimulated OV neurons while suppressing response gain in others. Other pairs with elevated *S*^+*−*^ involve the stimulated OV *−* group (OV *−* /offer-aligned CJ, OV*−*/CV+, OV*−*/CV*−*, and OV*−*/UC; green in Fig. 6BC). These pair types also show intermediate- to-high *S*^++^, indicating coupled activation and suppression responses between stimulated OV neurons and other tuning groups. Finally, the offer-aligned CJ/offer-aligned CJ and OV+/offer-aligned CJ group pairs (purple in Fig. 6B–D) show the clearest combination of strong co-activation and little opposite-sign recruitment or co-suppression (large *S*^++^ but small *S*^+*−*^ and *S*^*−−*^).

Together, these results show that the OV–CJ functional backbone operates within a richer pattern of neuronal recruitment than a simple activation relay. Stimulation of OV neurons engages a distributed pattern of coordinated activation and suppression within and across tuning groups, in which the corresponding choice population is preferentially recruited.

### Avalanche activation–suppression structure is spatially organized on a lattice

Since neurons with the same dominant tuning can exhibit opposite recruitment signs during the same avalanche response, we asked whether this heterogeneity reflects a systematic spatial organization rather than merely idiosyncratic differences among neurons. To test this, we analyzed joint recruitment as a function of stereotaxic *z*-separation, *d*_*z*_, between neurons on the same probe (Fig. 7A).

**Figure 7.**
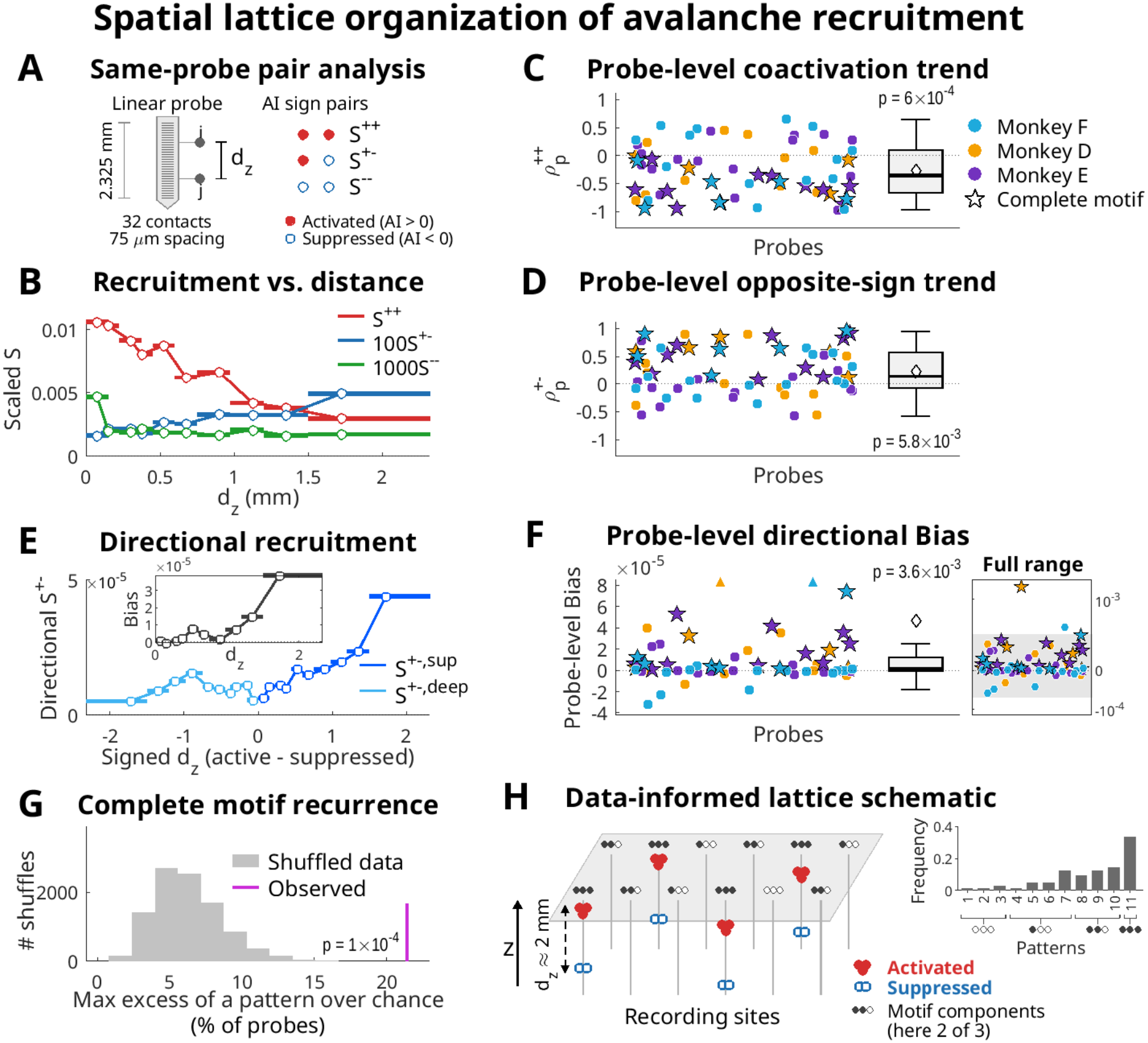
Spatial lattice organization of avalanche recruitment. **(A)** Schematic of same-probe neuron-pair analysis. (**B**) Mean same-probe Synchrony Indices across pooled neuron-pair observations in 10 global equal-count distance bins. Circles mark means at median *d*_*z*_; bars span bin ranges. *S*^+*−*^ and *S*^*−−*^ are scaled by 100 and 1000, respectively. Co-activation decreases and opposite-sign recruitment increases with distance, while co-suppression is enriched in the nearest bin. (**C**,**D**) Probe-level Spearman correlations of co-activation 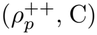 and opposite-sign recruitment 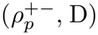 with *d*_*z*_ at 63 recording sites. Points represent sites, color-coded by animal; stars mark sites exhibiting the complete motif 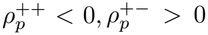, and directional Bias *B*_*p*_ *>* 0). Boxes span the 25th–75th percentiles; lines mark medians, diamonds means, and whiskers extend to the outermost data points within 1.5 IQR below or above the box. Across-site medians confirm decreasing co-activation and increasing opposite-sign recruitment with distance (*p* = 6.0*×* 10^*−*4^ and *p* = 0.0058, respectively; two-sided, Holm-adjusted spatial-permutation tests). (**E**) Mean opposite-sign recruitment with the active component superficial (*S*^+*−*, sup^) or deep (*S*^+*−*, deep^) plotted at +*d*_*z*_ and *−d*_*z*_, respectively, using the bins in (B). Inset: Bias (*S*^+*−*, sup^ *− S*^+*−*, deep^) versus *d*_*z*_. (**F**) Probe-level directional Bias, *B*_*p*_, in the largest-distance bin (1.5–2.325 mm). Plotting conventions as in (C,D). Triangles mark two Bias values above the main-panel range. Inset: the same 63 sites without clipping, on an asinh scale (approximately linear near zero, logarithmic at large magnitudes). Shading marks the main-panel range; the two clipped sites appear above it as a star (complete motif) and a circle. The median *B*_*p*_ confirms a systematic preference for activation superficial to suppression (*p* = 0.0036; tested as in C,D). (**G**) Enrichment of the complete motif in the observed data (magenta), compared with the maximum enrichment across all 8 sign patterns in each of 10,000 spatial shuffles (gray). Each pattern is one combination of positive or negative signs for 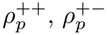, and *B*_*p*_. Enrichment is the difference between a pattern’s observed frequency and its mean shuffled frequency. No shuffled maximum reached the observed complete-motif enrichment (*p* = 1.0 *×*10^*−*4^). (**H**) Data-informed lattice schematic. “Lattice” denotes the preferential recurrence across recording sites of a characteristic motif of activation (red) and suppression (blue). The complete motif is drawn only at sites sharing all 3 defining relationships (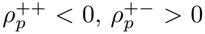, *B*_*p*_ *>* 0). Filled gray dots indicate how many relationships each site’s pattern shares with the complete motif (0–3); sites sharing each number of relationships are represented in proportions approximating those in the data (top right). Top right: frequencies of all 11 patterns identified in the data (8 from (G) and 3 with 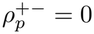, *B*_*p*_ = 0, or both).

Same-probe pairs were pooled across recording sites, sessions, and offer simulations to define 10 equal-count (quantile) distance bins, within which we computed mean *S*^++^, *S*^+*−*^, and *S*^*−−*^. The pooled and probe-level analyses below reveal a spatial lattice organization of avalanche recruitment. We use “lattice” in a statistical sense to denote the significantly enriched recurrence across recording sites of a characteristic spatial motif of activation and suppression, without requiring observation of the complete motif at every site or registration to fixed absolute cortical depths. The supporting relationships are quantified in Fig. 7B–G and summarized in Fig. 7H.

Fig. 7B shows the mean Synchrony profiles across distance bins. Co-activation (*S*^++^) is 3.57-fold greater in the nearest-distance bin than in the largest-separation bin and decreases with *d*_*z*_ (Spearman *ρ*^++^ = 0.976; 95% session-bootstrap CI [*−*0.988, *−*0.758]). In parallel, opposite-sign recruitment (*S*^+*−*^) increases with *d*_*z*_ (*ρ*^+*−*^ =*−* 0.903; 95% CI [0.600, 0.988]), reaching a 3.01-fold higher level in the largest-separation bin than in the nearest bin. Its maximum at 1.5–2.325 mm corresponds to an approximate 2 mm spacing between activation and suppression. Together, these pooled profiles reveal substantial spatial modulation linking local co-activation to spatially offset suppression. Co-suppression (*S*^*−−*^) is enriched by a factor of 2.49 in the nearest-distance bin (*d*_*z*_*≤*0.15 mm) relative to all remaining pairs (95% session-bootstrap CI [1.36, 3.98]), indicating local clustering of co-suppressed neurons.

The different numerical scales of the three Synchrony Indices reflect the AI magnitudes they multiply. For neurons not directly stimulated, |AI| = 0.01 means that, at the steepest point of their response, firing changes by 2 spikes/s per unit increase in offer input. At low baseline firing rates, neurons have little activity to suppress before reaching zero, whereas activation can recruit substantial additional activity (Fig. 1E). The smaller scales of *S*^+*−*^ and *S*^*−−*^ are consistent with this physiological constraint. Nevertheless, all three recruitment modes exhibit pronounced spatial modulation.

We next tested whether decreasing co-activation and increasing opposite-sign recruitment with distance recur across recording sites (Fig. 7CD). For this and subsequent probe-level tests, we generated a spatial null using 10,000 permutations that randomly reassigned each neuron’s offer-specific Avalanche Indices together across spatial positions within each session and probe (Methods). This preserved the joint distribution of responses across offers and the sampling geometry while disrupting the association between recruitment and position. With fewer neuron pairs per distance bin, individual probe profiles are more susceptible to sampling fluctuations that can weaken rank correlations relative to the pooled profile. Nevertheless, probe-level Spearman correlations of *S*^++^ and *S*^+*−*^ with *d*_*z*_ confirm a systematic tendency across recording sites toward decreasing co-activation (median 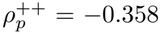; IQR [*−*0.671, 0.091]; *p* = 6.0 *×*10^*−*4^) and increasing opposite-sign recruitment (median 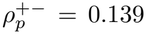; IQR [*−* 0.073, 0.570]; *p* = 0.0058; two-sided tests of across-site medians against the spatial null, Holm-adjusted).

This activation–suppression motif has a preferred orientation along the cortical depth axis, with its active component located at a shallower depth than its suppressed component. To quantify this directional preference, we defined the signed offset *z*_active_ *− z*_suppressed_ for each same-probe neuron pair with opposite-sign responses and assigned its *S*^+*−*^ contribution to *S*^+*−*, sup^ for positive offsets (activation superficial to suppression) or to *S*^+*−*, deep^ for negative offsets (activation deeper than suppression; Methods). Fig. 7E displays the distance bins from Fig. 7B on a signed axis, with mean *S*^+*−*, sup^ plotted at positive separations and mean *S*^+*−*, deep^ at negative separations. The inset shows the directional Bias, defined in each absolute-distance bin as the difference between the mean *S*^+*−*, sup^ and *S*^+*−*, deep^. Active-superficial recruitment exceeds active-deep recruitment in nine of the ten bins, with the greatest Bias in the largest-distance bin (1.5–2.325 mm), where *S*^+*−*^ is also maximal. Fig. 7F shows the directional Bias, *B*_*p*_, computed separately for each probe within this bin. Probe-level Bias confirms a systematic preference for activation superficial to suppression across recording sites (median *B*_*p*_ = 1.53*×*10^*−*6^; interquartile interval [*−* 0.738, 12.3] *×*10^*−*6^; *p* = 0.0036; two-sided, Holm-adjusted permutation test against the spatial null).

Finally, we asked whether the three spatial relationships – decreasing co-activation with distance, increasing opposite-sign recruitment with distance, and positive directional Bias – converge within the same recording sites (Fig. 7G). We considered all 8 combinations of positive and negative values of the two distance correlations (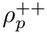 and 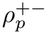) and probe-level directional Bias (*B*_*p*_). For each of the 8 patterns, we calculated its expected frequency under the spatial null by averaging the proportion of sites exhibiting that pattern across the permutations. We then quantified each pattern’s enrichment as its observed frequency minus its mean frequency under the spatial null. We compared the largest observed enrichment across patterns with the largest enrichment across patterns in each permutation (Methods). The complete motif (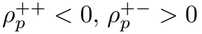, and *B*_*p*_ *>* 0) is both the most frequent observed pattern and the most enriched of the 8 patterns, occurring at 21*/*63 sites (33.3%), compared with 11.9% under spatial shuffling, approximately 2.8-fold the shuffled expectation. None of the 10,000 permutations produces an equally large or larger enrichment for any pattern (*p* = 1.0 *×*10^*−*4^). Of the remaining sites, 23*/*63 (36.5%) exhibit patterns sharing two of the three defining relationships of the complete motif, 15*/*63 (23.8%) share one, and 4*/*63 (6.3%) share none (Fig. 7H, frequency plot). Additionally, the complete motif is the most frequent and most enriched pattern in each monkey, occurring at 5*/*16 sites in D (31.3%), 10*/*26 in E (38.5%), and 6*/*21 in F (28.6%).

### Neuron-resolved interaction structure organizes lattice-patterned recruitment beyond dominant tuning

Because different tuning-pair types may occur at different frequencies across *d*_*z*_ and differ in their mean *S*^++^ and *S*^+*−*^ values (Fig. 6BC), we asked whether the spatial dependence simply reflects tuning composition or persists after controlling for it. For example, if aligned-CJ/aligned-CJ pairs, which have relatively high mean *S*^++^, were more frequent at short *d*_*z*_, while OV*−* /CV+ pairs, which have relatively high mean *S*^+*−*^, were more frequent at large *d*_*z*_, this alone could produce decreasing *S*^++^ and increasing *S*^+*−*^ with distance. Alternatively, even within the same tuning-pair type, neuron pairs could exhibit stronger co-activation at short distances and stronger opposite-sign recruitment at larger distances, revealing spatial organization beyond tuning composition. To test this, we computed tuning-corrected *S*^++^ and *S*^+*−*^ that remove differences in mean values across tuning-pair types while preserving variation within each type (Methods). Both pooled distance trends remain strong after correction: co-activation decreases with *d*_*z*_ (*ρ*^++^ = *−*0.939; 95% session-bootstrap CI [*−*0.976, *−*0.576]), and opposite-sign recruitment increases (*ρ*^+*−*^ = 0.915; 95% CI [0.661, 0.976]; Fig. S2AB). All three relationships also persist across recording sites (median 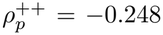, *p* = 0.0012; median 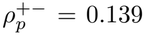, *p* = 0.024; median *B*_*p*_ = 1.84 *×* 10^*−*6^, *p* = 0.0052; two-sided, Holm-adjusted spatial-permutation tests; Fig. S2CDE). The complete motif remains the most enriched sign pattern, occurring at 22.2% of sites versus 11.1% under spatial shuffling (*p* = 0.037; permutation test; Fig. S2F). Thus, the lattice is not simply a by-product of the spatial distribution of dominant tuning.

We next asked if this recruitment organization could nevertheless emerge from coarse interactions between tuning groups. To test this, we replaced neuron-specific interactions with tuning-group means. Although retaining neuron-specific biases *h*_*i*_ could in principle preserve heterogeneous recruitment within groups, the systematic subdivision into activated and suppressed neurons is lost, and neither the three probe-level lattice components nor the complete motif differs significantly from the spatial null (Fig. S3). Thus, coarse tuning-group interactions capture the aggregate transformation, whereas finer-grained interactions shape its network-specific expression through patterned activation, suppression, and spatial lattice organization.

## Discussion

By perturbing data-inferred OFC networks, we identified a network-level mechanism that transforms offer values into choice-selective attractor states. The inferred interactions coordinate avalanche responses, biasing collective recruitment toward the choice population favored by relative value (Figs. 1–3). Targeted ablations (Fig. 4), network reductions (Fig. 5), and neuron-resolved analyses (Figs. 6–7) reveal the functional interaction pathways that drive the transformation and how the resulting neuronal recruitment is systematically organized.

The ablations identify OV–CJ interactions as the backbone of the transformation: removing them virtually abolishes choice bias, whereas retaining only the OV–CJ core preserves approximately 60% of the full-network bias (Fig. 4). The broader network, including CV and UC populations, further amplifies this effect. Both negative- and positive-encoding pathways independently support the transformation, with the latter contributing more strongly. Their effects largely add, providing complementary routes from value to choice. Thus, while previous studies identified population-level signatures consistent with attractor dynamics [21, 22, 34–39], our results identify the interaction-level components through which choice-selective attractor states emerge in inferred OFC networks.

In parallel, our mean-field reductions reveal that mean interactions between tuning groups preserve the direction and 51–68% of the magnitude of the transformation on average across networks and input conditions, but poorly reproduce the choice bias of individual networks under specific inputs (Fig. 5). Thus, coarse interactions capture much of the transformation at the aggregate level, whereas finer structure is needed to reproduce its network- and condition-specific realization. One manifestation of this finer structure is the splitting of similarly tuned neurons into activated and suppressed subpopulations during avalanches (Fig. 6). These contrasting responses suggest complementary computational roles: activation may strengthen pathways supporting the offer-aligned choice representation, while suppression may weaken influences opposing its recruitment or supporting the competing choice. Their coordination within tuning groups could provide fine-grained control over signal propagation, enhancing its selectivity and sharpening choice-selective recruitment.

A statistical lattice organizes these recruitment roles in space: co-active neurons tend to lie close together, while activation and suppression are preferentially separated by approximately 2 mm along the probe axis, with suppression deeper below the cortical surface (Fig. 7). This lattice suggests a reusable spatial architecture underlying offer-dependent activation and suppression: different offers preferentially recruit their corresponding choice populations within a common spatial scaffold. In particular, the preferred orientation of this motif is consistent with descending inhibitory circuits described in other cortical areas [58–60]. This lattice organization is not reducible to the spatial distribution of tuning groups (Fig. S2). Both within-group splitting and the spatial lattice are lost when neuron-specific interactions are replaced with tuning-group means, indicating that finer interactions organize recruitment within functional groups and across space (Fig. S3).

Several features of our modeling framework are important for interpreting these results. First, in the full inferred networks, neurons were classified by dominant tuning, but these labels served only to define controlled inputs, specify targeted ablations, and summarize network responses. Interactions were inferred directly from population activity without imposing any tuning-defined structure. Thus, the observed attractor transitions do not require separate neuronal groups with single-variable selectivity. We treat tuning labels as convenient projections for probing the network, not as fundamental building blocks of the computation. The mean-field reductions are a deliberate exception: they explicitly replace neuron-specific interactions with tuning-group averages to identify which properties of the transformation are preserved by this reduction and which require finer structure. Second, our model is fully constrained by observed firing rates and pairwise correlations. Thus, the observed attractor transitions do not require irreducible higher-order interactions. Third, the inferred Ising couplings represent effective interactions within an equilibrium model whose states can be sampled through Monte Carlo simulations and whose responses to inputs can be probed through susceptibility. These interactions can reflect both direct and indirect influences mediated by unrecorded neurons. Our results therefore show that the effective interaction structure expressed in OFC is sufficient to produce the value-to-choice transformation, without requiring the computation to be confined to OFC. Related computations may occur across interconnected regions [61], whose inputs may contribute to the effective interactions observed in OFC.

In conclusion, inferred pairwise effective interactions in OFC transform offer values into choice-selective attractor states through an OV–CJ backbone, amplified by the broader network. Interaction structure beyond tuning-group averages shapes the network-specific realization of the transformation, coupling neuronal activation and suppression into a recurring spatial motif.

## Methods

### Experimental procedures

All procedures conformed to the NIH Guide for the Care and Use of Laboratory Animals and were approved by the Institutional Animal Care and Use Committee at Washington University. The experimental protocol is described in full detail in [55] and summarized below.

Three adult male rhesus monkeys (*Macaca mulatta*) performed a binary juice-choice task. In each session, animals chose between variable quantities of two juices, A and B (with A preferred), presented simultaneously on opposite sides of fixation. Juice identity and quantity were indicated by the color and number of visual symbols, respectively, and choices were reported by saccade. Offer pairs were presented pseudorandomly, with left–right position counterbalanced across trials. Sessions included 181–402 trials (median = 264) and lasted 30–60 min; across sessions, 12 juice types and 19 juice pairings were used.

For each session, we estimated the relative value of the two juices, *ρ*, from the animal’s choices using a logistic regression as a function of log(*q*_*B*_*/q*_*A*_), where *q*_*A*_ and *q*_*B*_ are the quantities of juices A and B offered on each trial. The parameter *ρ* corresponds to the indifference point of the fitted logistic, at which the animal splits its choices equally between the two options.

Extracellular recordings targeted the orbital gyrus (area 13m) and were obtained across 85 sessions from both hemispheres in two monkeys and from the right hemisphere in the third one. Recording locations were guided by structural MRI. Neural activity was recorded with 32-channel linear probes (V-probe, Plexon; 120 *µ*m diameter) with contacts spaced by 75 *µ*m. One to four probes were used per session, with at most two probes per hemisphere; probes were advanced with motorized microdrives at 1 *µ*m resolution. Spikes were identified and sorted offline.

Before performing the computational analyses, we defined objective session inclusion criteria to ensure that the inferred networks could be meaningfully simulated and compared across offer conditions. Specifically, we included sessions in which at least 7 of the 9 neuron groups defined by dominant tuning were sampled. In addition, simulations under both offer conditions required at least one corresponding input neuron for each offer type (i.e., at least one OVA+ or OVB*−* neuron and at least one OVB+ or OVA *−*neuron). Finally, both CJA and CJB neurons had to be present to enable readout of choice preference from the inferred networks. Applying these criteria reduced the initial dataset from 85 to 63 sessions, all of which were included in the Monte Carlo simulations described below. As detailed in the Monte Carlo convergence analysis, one session did not converge for the single-offer simulations and was excluded from all subsequent simulation analyses. A second session did not converge for the competing-offer simulations and was excluded from those analyses only.

### Easy- and hard-trial classification

For each trial, we computed a normalized value-contrast index,

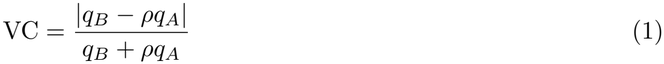

The index quantifies the relative distance of the offered values from the session-specific indifference condition, *q*_*B*_ = *ρq*_*A*_. It equals 0 at indifference and increases as the relative imbalance between the two offer values increases, reaching its maximum of 1 when only one offer is present.

In Fig. 3B, within each session, trials were divided according to the median VC. Trials with VC above the median were classified as easy, whereas trials with VC below the median were classified as hard. Trials with VC exactly equal to the median were excluded.

### Ising model and network inference

For each recording session, we modeled the simultaneously recorded neuronal population using an Ising model. Each neuron *i* was represented by a binary variable *σ*_*i*_ *∈ {*0, 1*}*, indicating whether the neuron was silent or active in a given 5-ms time bin. The model assigns the following probability to each population activity pattern ***σ*** = (*σ*_1_, … , *σ*_*N*_):

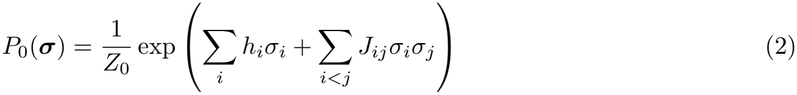

where *h*_*i*_ represents the intrinsic bias of neuron *i* toward activity, *J*_*ij*_ represents the effective pairwise interaction between neurons *i* and *j*, and *Z*_0_ is the partition function ensuring normalization. The energy of the model in the absence of external inputs is

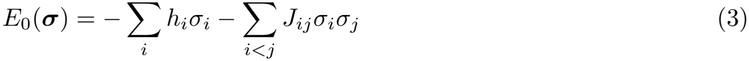

such that 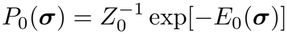. The interactions are symmetric (*J*_*ij*_ = *J*_*ji*_), with no self-interactions (*J*_*ii*_ = 0). Model parameters were inferred using the Adaptive Cluster Expansion (ACE) algorithm, previously developed in [46–49], which estimates the biases *h*_*i*_ and interactions *J*_*ij*_ required to reproduce the empirical firing rates and pairwise correlations measured in 5-ms time bins. Among distributions reproducing these statistics, the Ising model has maximum entropy and is therefore the least biased distribution consistent with the measured constraints. In all analyses except that shown in Fig. 3B, a single model was inferred for each recording session using all trials, spanning different offer quantities and choices. Activity patterns spanning trial onset to trial offset were concatenated across trials for model inference. In Fig. 3B, separate Ising models were inferred from the easy- and hard-trial subsets of each session.

### Monte Carlo simulations of the inferred Ising models

We sampled population activity states from the inferred Ising models under external inputs using the Metropolis–Hastings algorithm [56, 57]. External inputs **H** = (*H*_1_, … , *H*_*N*_) were applied during these simulations to probe the network response, while the inferred parameters *h*_*i*_ and *J*_*ij*_ were kept fixed. The external inputs modify the energy of the inferred model from *E*_0_(***σ***) to

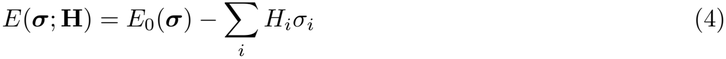

and therefore define the corresponding equilibrium Ising distribution

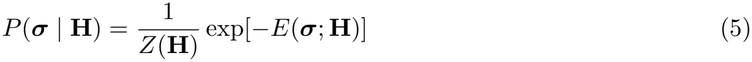

where *Z*(**H**) = ∑_***σ***_ exp[*−E*(***σ***; **H**)] is the partition function. For **H** = 0, this distribution reduces to the originally inferred model, *P*_0_(***σ***).

The Metropolis–Hastings algorithm evolves the network through successive single-neuron state updates. A single Monte Carlo (MC) flip attempt consists of proposing a change in the state of one neuron, *σ*_*i*_ *→* 1 *− σ*_*i*_, and accepting the proposed state with probability min[1, exp(*−*Δ*E*)], where Δ*E* is the resulting change in energy. One MC sweep is defined as *N* such flip attempts, where *N* is the number of neurons in the network. By iterating this procedure over many MC sweeps, the Markov chain approaches stationarity, allowing network states to be sampled from the equilibrium distribution *P* (***σ*** | **H**) of the model.

Before production sampling, each simulation was run through successive pre-run windows to assess equilibration. Convergence was evaluated from the mean-activity and susceptibility estimates defined below by asking whether changes between successive windows were larger than expected from Monte Carlo sampling uncertainty, whether unusually large discrepancies persisted in the independent production run, and whether estimates showed systematic drift. We also examined the prevalence of extreme discrepancies within each session to identify session-specific convergence failures. The full convergence procedure and results are provided in the Supplementary Information. After this pre-run convergence assessment, states were recorded every 5 MC sweeps to reduce correlations between successive samples, and 20,000 production samples were collected for each simulation to estimate neuronal mean activities and susceptibilities. In each simulation, we applied a uniform baseline input *H*_base_ to all neurons to place the network in a responsive regime in which selective offer inputs could reveal coordinated population responses supported by the inferred interaction structure. The baseline input was chosen such that 10% of neurons had a mean activity *⟨σ*_*i*_*⟩ ≥* 0.1. The total external input to each neuron was defined as

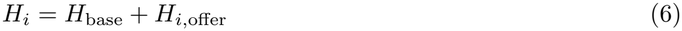

where *H*_*i*,offer_ denotes the offer-specific inp ut. We define the two offer-aligned input populations as

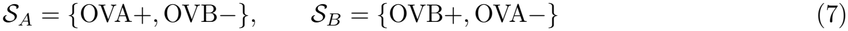

For the single-offer A simulations, we set *H*_*i*,offer_ = *H*_*A*_ *>* 0 for neurons *i ∈ S*_*A*_, assigned a sufficiently large negative input to neurons *i ∈ S*_*B*_ to keep their activity at zero, and set *H*_*i*,offer_ = 0 for all other neurons. Conversely, for the single-offer B simulations, we set *H*_*i*,offer_ = *H*_*B*_ *>* 0 for neurons *i ∈ S*_*B*_, assigned the negative silencing input to neurons *i ∈ S*_*A*_, and set *H*_*i*,offer_ = 0 otherwise (Fig. 1C). For the competing-offer simulations, we set *H*_*i*,offer_ = *H*_*A*_ for neurons *i ∈ S*_*A*_, *H*_*i*,offer_ = *H*_*B*_ for neurons *i ∈ S*_*B*_, and *H*_*i*,offer_ = 0 otherwise. In each single-offer simulation, only the positive offer-specific input amplitude, *H*_*A*_ or *H*_*B*_, was systematically varied, whereas *H*_base_ and the negative silencing input were held fixed. In competing-offer simulations, both positive offer inputs were varied together at a fixed ratio, as detailed below. Neuronal mean activities, defined over 5-ms time bins, and susceptibilities were computed from the sampled states at each input level. For simplicity, we refer to neurons receiving the variable positive offer input as stimulated, and to all other neurons as non-stimulated.

### Neuronal mean activities and susceptibilities

We denote equilibrium averages at fixed **H** by *⟨·⟩*. The mean activity of neuron *i* is

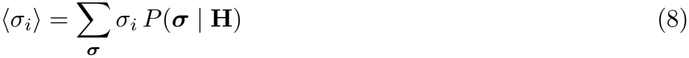

In practice, this expectation was estimated from the *M* = 20,000 network states sampled in the MC simulation after equilibration using the sample mean,

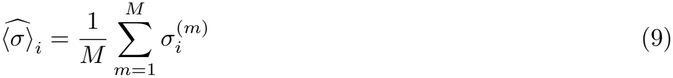

where 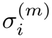 denotes the state of neuron *i* in the *m*th sampled network state.

The susceptibility of neuron *i* to an external input applied to neuron *j* is defined as

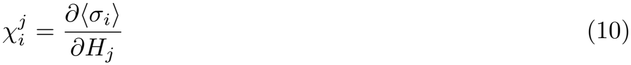

In principle, this derivative could be estimated by comparing mean activities across simulations with nearby input values. However, finite-difference estimates are sensitive to sampling noise. We therefore used an equivalent formulation that expresses susceptibilities in terms of covariances between neuronal activities at fixed **H**. Using 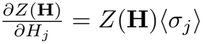, we obtain:

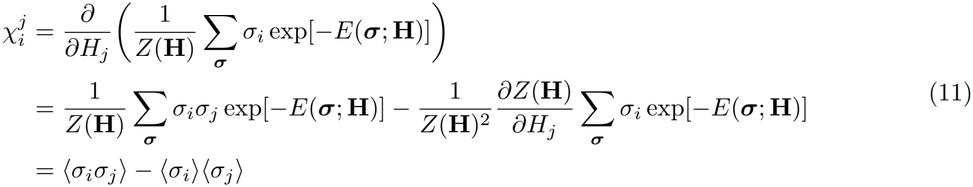

Thus, the susceptibility of neuron *i* to the external input applied to neuron *j* is equal to the covariance between their activities.

#### Neuronal susceptibilities to single-offer inputs

Let *X ∈ {A, B}* label the single-offer simulation condition. When the same positive offer-specific input amplitude *H*_*X*_ is applied to a subset of neurons *S*_*X*_ , while the negative silencing input and *H*_base_ are held fixed, only the inputs to neurons in *S*_*X*_ vary with *H*_*X*_. The susceptibility of neuron *i* is therefore:

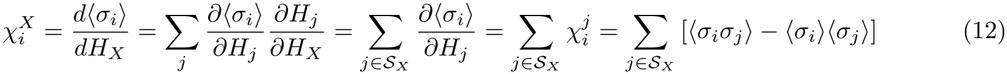

For each simulation condition, susceptibilities were estimated directly from the *M* = 20,000 sampled network states by replacing the equilibrium averages in Eq. (12) with their sample estimates:

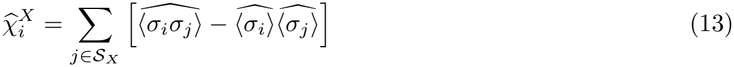

#### Neuronal susceptibilities to simultaneous competing-offer inputs

To generate the results in Fig. 3, we simultaneously applied stimulation *H*_*A*_ = *cH*_*A*0_ to *S*_*A*_ neurons, and stimulation *H*_*B*_ = *cH*_*B*0_ to *S*_*B*_ neurons. *H*_*A*0_ and *H*_*B*0_, and hence the ratio

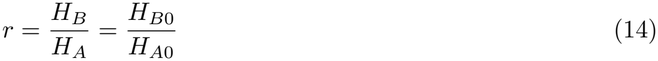

were held fixed for each input-bias condition, whereas *c*, and therefore the total input

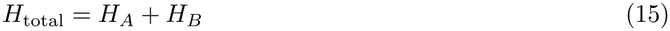

were varied. Here, *c* serves as a control parameter that probes the internal susceptibility of the network to offer inputs by scaling the overall influence (or salience) of the inputs, whereas *r* represents the input bias toward offer B.

By applying the same drive *H*_*A*_ to all neurons in *S*_*A*_ and the same drive *H*_*B*_ to all neurons in *S*_*B*_, we simulated conditions in which the two offers occupied approximately complementary positions within their respective value ranges, i.e., 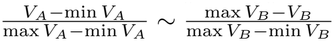

The susceptibility of neuron *i* was defined as the change in its equilibrium mean activity with respect to *H*_total_ along trajectories of fixed *r*. Since

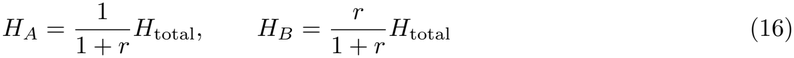

and *H*_base_ is held fixed, the chain rule and Eq. (11) give

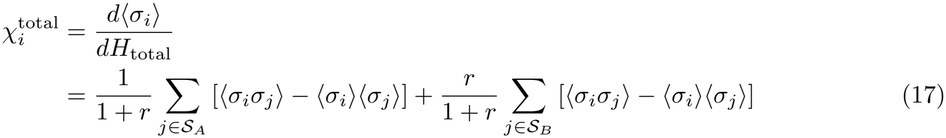

At each value of *H*_total_ and fixed *r*, 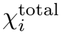 was estimated from the *M* = 20,000 sampled network states by replacing the equilibrium averages in Eq. (17) with their corresponding sample estimates.

### Susceptibility peaks and Avalanche Index

For each neuron *i* and input type *X ∈ {A, B*, total*}*, we identified the positive input strength *H*_*X*_ at which the absolute magnitude of the estimated susceptibility was maximal. Here, *X* = *A* and *X* = *B* refer to the corresponding single-offer inputs, whereas *X* = total refers to the competing-offer input *H*_total_ = *H*_*A*_ + *H*_*B*_, varied at fixed input ratio *r* = *H*_*B*_*/H*_*A*_. Thus,

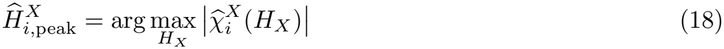

For each simulation condition, susceptibility peaks were identified over the discrete grid of input strengths used in the Monte Carlo simulations. For single offers, this grid was *{*0, 0.3, … , 9.9*}*. For competing offers, *H*_*A*_ = *{*0, 0.3, … , 9.9*}* and *H*_*B*_ = *r × {*0, 0.3, … , 9.9*}* if *r ≤* 1; *H*_*B*_ = *{*0, 0.3, … , 9.9*}* and 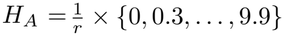 if *r >* 1.

We then obtained the signed peak susceptibility, corresponding to either a maximum or a minimum, as:

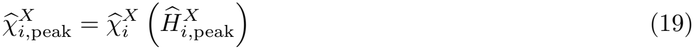

For neurons receiving a variable positive offer input, part of the susceptibility arises directly from the external input, even in the absence of network interactions. To treat single and competing offers uniformly, we define

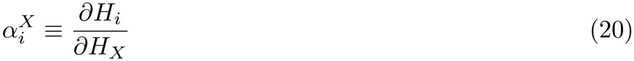

For the single-offer conditions,

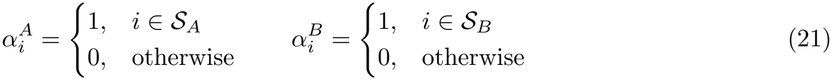

where the negative silencing inputs are held fixed as *H*_*A*_ or *H*_*B*_ is varied. For competing offers,

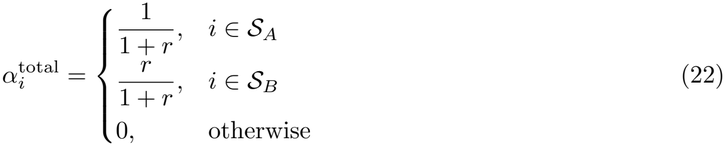

For an isolated binary neuron,

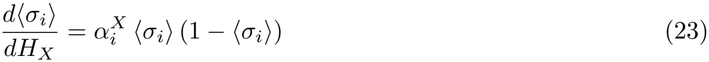

whose maximal value is 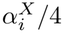. We therefore define the baseline peak susceptibility, corresponding to the maximal direct susceptibility expected in the absence of network interactions, as

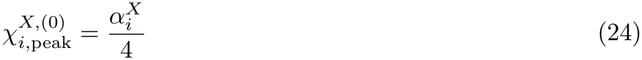

The Avalanche Index was then defined as

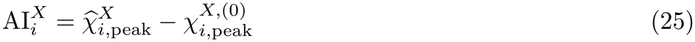

The Avalanche Index therefore quantifies the signed maximal rate of change of each neuron’s activity with the offer input, after subtracting the maximal direct contribution expected in the absence of network interactions.

### Susceptibility-peak coordination analyses

To determine whether the inferred network interactions promote the alignment of neuronal susceptibility peaks, we examined the input strengths at which neurons reached their maximal absolute susceptibility during single-offer simulations (Fig. 1F). We performed two complementary analyses. First, because directly stimulated OV neurons exhibit susceptibility peaks even without interactions, we tested whether recurrent interactions reduce the dispersion of their peak locations relative to an independent-neuron baseline. Second, because non-stimulated neurons have zero susceptibility in the absence of interactions, we assessed whether their network-mediated susceptibility peaks preferentially align with those of the stimulated OV population of their own network compared with those of other networks.

#### Coordination among stimulated OV neurons

For each session *s* and offer *X ∈ {A, B}*, we considered the directly stimulated OV population *S*_*X*_ and quantified the dispersion of its susceptibility-peak locations as

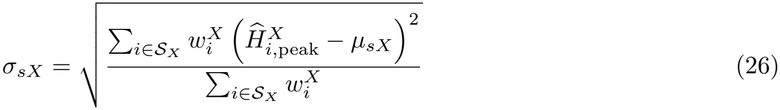

Where

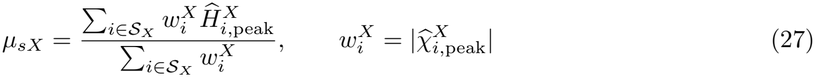

Smaller *σ*_*sX*_ indicates tighter alignment of susceptibility peaks among stimulated neurons. Weighting by absolute peak susceptibility gives greater importance to neurons with stronger responsiveness.

We compared this dispersion with the prediction for independent neurons, obtained by setting all pairwise interactions to zero (*J*_*ij*_ = 0) while preserving each neuron’s intrinsic bias and external inputs. From Eq. (23), the susceptibility of an independent, directly stimulated neuron 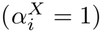 reaches its maximum when *⟨σ*_*i*_*⟩* = 1*/*2, corresponding to

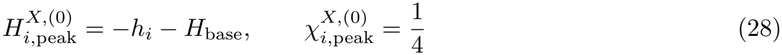

These expressions give the analytical maxima for unconstrained input strengths. For consistency with the inferred network simulations, we determined the independent-neuron peak locations and corresponding susceptibilities on the same discrete input grid (*H*_*X*_ = 0, 0.3, … , 9.9). We then computed the independent-neuron dispersion 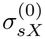 using the same weighted standard deviation as for the inferred networks, with the peak center and weights calculated from the independent-neuron peak locations and susceptibilities.

For each session, we computed the dispersions separately for offers A and B, provided that at least two directly stimulated OV neurons were available for that offer, and then averaged the results across eligible offers. In Fig. 1F, left, we plotted the resulting session-level dispersions, *σ*_*s*_ versus 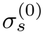, for the inferred networks and independent neurons, respectively, and compared them using a two-sided Wilcoxon signed-rank test. We also calculated the median paired difference 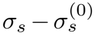 and its 95% confidence interval using 10,000 bootstrap resamples of sessions.

As sensitivity analyses, we repeated the comparison using (a) the peak susceptibility magnitudes from the inferred networks as weights for both inferred-network and independent-neuron peak locations, or (b) uniform weights for all neurons. The reduction in peak dispersion in the inferred networks remained significant under both alternative weighting schemes (*p* = 0.0032 and *p* = 0.0068, respectively).

#### Coordination of propagated responses

We next tested whether the susceptibility peaks of neurons not receiving the variable positive offer input were preferentially aligned with the stimulated OV population of their own network. For each session *s* and offer *X ∈ {A, B}*, we defined the stimulated OV peak center *µ*_*sX*_ as above and calculated the susceptibility-weighted mean absolute distance of the remaining neurons’ peak locations from this center:

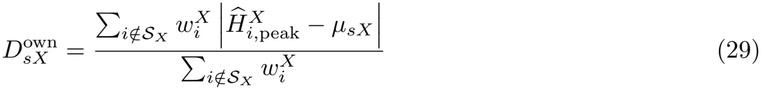

Smaller values indicate tighter alignment of non-stimulated neurons’ susceptibility peaks with the peak center of the stimulated OV population.

To assess whether this alignment was specific to each network, we generated *K* = 10,000 random reassignments of stimulated OV peak centers across sessions, each constrained so that no session retained its own center (derangements). The susceptibility-peak locations and weights of non-stimulated neurons remained unchanged, and the same reassignment was applied to both offers, preserving their pairing within sessions. For each session *s*, offer *X*, and permutation *b*, we calculated

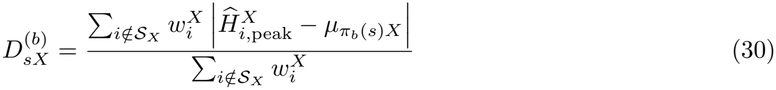

where *π*_*b*_(*s*)*≠s* denotes the session assigned to session *s* in permutation *b*.

For each permutation and offer, we averaged these distances across sessions to construct a null distribution, against which we compared the mean own-network distance, 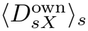. Two-sided Monte Carlo permutation *p*-values were calculated using the plus-one correction, with Holm adjustment across the two offers.

To quantify the coordination effect for each session and offer, we calculated the mean mismatched distance

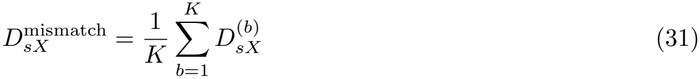

and defined

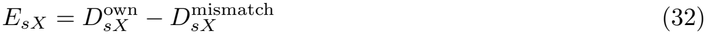

Fig. 1F, right, plots 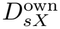 versus 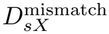. Negative values of *E*_*sX*_ indicate closer alignment with the own-network OV center than with the OV centers of other networks. We calculated the median coordination effect and its 95% confidence interval using 10,000 bootstrap resamples of sessions, separately for each offer.

For each offer, the percentage reduction in mean distance relative to mismatched network centers was calculated as 100 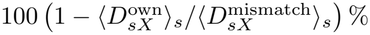.

#### Susceptibility Peak Alignment

For Fig. 2C, we quantified the alignment of susceptibility peaks across all non-silenced neurons in each single-offer simulation. For each session *s* and offer *X ∈ {A, B}*, we defined the Alignment as the inverse susceptibility-weighted peak dispersion:

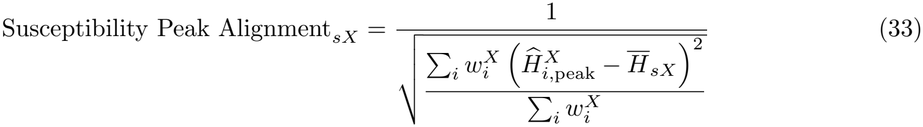

Where

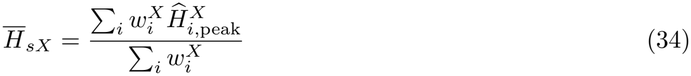

Here, the sums extend over all non-silenced neurons in the network, with weights 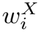 defined above. We compared Alignment between conditions with elevated and non-elevated mean Avalanche Index of CJ neurons (defined below) using a two-sided Wilcoxon rank-sum test.

#### Avalanche Asymmetry for single and competing offers

To quantify whether avalanche recruitment preferentially favored one choice population over the other, we computed Avalanche Asymmetry from the neuron-specific Avalanche Indices. For any neuronal group *g* and session *s*, its mean Avalanche Index under input type *X* is

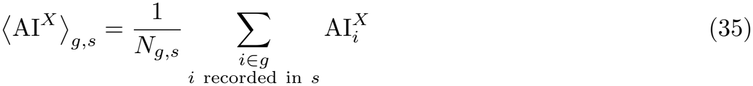

where *N*_*g,s*_ is the number of neurons in group *g* recorded in session *s*.

For each session *s*, the single-offer Avalanche Asymmetry was defined as the difference between the mean AI of the offer-aligned and offer-opposed choice populations:

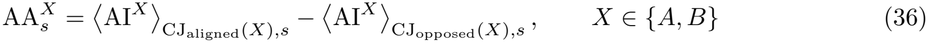

Thus,

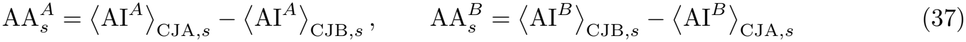

Positive values therefore indicate preferential recruitment of the choice population aligned with the simulated offer. We refer to this quantity as offer-aligned Avalanche Asymmetry. This convention was used when the two single-offer conditions were summarized relative to the simulated offer, including the ablation analyses in Fig. 4.

For the competing-offer simulations, the sign convention was instead held fixed as CJB*−*CJA across input configurations. For session *s*, we write 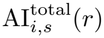 for the Avalanche Index obtained by varying *H*_total_ at fixed *r* = *H*_*B*_*/H*_*A*_. Avalanche Asymmetry was then defined as

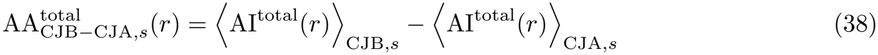

Positive values indicate stronger recruitment of CJB neurons, negative values indicate stronger recruitment of CJA neurons, and zero indicates equal recruitment of the two choice populations. The fixed CJB *−* CJA convention therefore allows Avalanche Asymmetry to track the direction and magnitude of each session’s choice bias as the competing-offer input bias *r* is varied. At *r* = 1 it quantifies any intrinsic B-versus-A choice bias of the inferred network under symmetric competing input.

For analyses spanning both single- and competing-offer conditions (Figs. 3 and 5), we expressed Avalanche Asymmetry in a common fixed CJB *−* CJA coordinate. Let *κ* denote the input configuration, with *κ* = *A* and *κ* = *B* denoting the two single-offer conditions and *κ* = (total, *r*) a competing-offer condition. We define

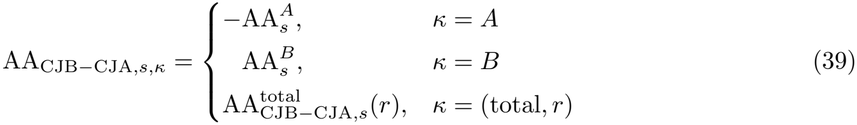

In this coordinate, negative values indicate A-directed recruitment and positive values indicate B-directed recruitment irrespective of stimulation protocol.

### Random network controls

For each inferred network and offer simulation, we generated 10 random network controls with matched statistics. Each random network contained the same number of neurons as the non-silent population in the corresponding inferred network and simulation condition, and the number of stimulated neurons was also matched. The pairwise interactions *J*_*ij*_ and neuron biases *h*_*i*_ of each random network were drawn from Gaussian distributions whose means and variances matched those of the corresponding *J*_*ij*_ and *h*_*i*_ distributions in the inferred network. We simulated the random networks using the same Monte Carlo procedure as for the inferred networks and computed susceptibility curves for all neurons.

For the analysis in Fig. S1, we calculated susceptibility-weighted peak dispersion separately for each of the 10 matched random-network realizations, using the same definition as for the inferred networks. For each session–offer condition, we defined random-network Alignment as the inverse of the mean peak dispersion across realizations. We compared inferred- and random-network Alignment across session–offer conditions using a two-sided Wilcoxon signed-rank test.

For each session–offer condition, we calculated the mean Avalanche Index, AI_rand_, by averaging across non-stimulated neurons and the 10 matched random realizations. The corresponding standard deviation, SD_rand_, was obtained by first calculating the standard deviation across non-stimulated neurons in each realization and then averaging over realizations. The mean AI of stimulated neurons was calculated similarly, pooling across the same realizations. Fig. 2A shows the mean*±* SEM of these values across sessions and both offer simulations (A and B).

Denoting by AI (CJ) the mean AI across the *N*_CJ_ neurons of the CJ population in the inferred network under a given simulation condition, we classified AI (CJ) as elevated relative to the matched random networks if it exceeded the random-network mean for non-stimulated neurons by more than three SEMs:

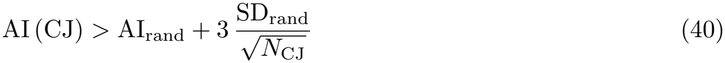

For Fig. 2D, we generated randomly connected networks using the same procedure, but without matching their parameters to the inferred networks. Instead, we systematically varied the mean of the *J*_*ij*_ distribution, the total number of neurons, and the number of stimulated neurons. For each parameter combination, we generated 50 random network realizations. The mean Avalanche Index was then computed by averaging across all non-stimulated neurons and across all realizations with the same parameters that satisfied the Monte Carlo convergence criterion (Supplementary Information).

For the ablation analyses in Fig. 4, we generated 10 identical replicas of each inferred network and independently ablated a random set of connections in each replica, matching the number of connections removed in the corresponding targeted ablation of the inferred network. Avalanche Asymmetry was averaged across the 10 random realizations for each session–offer c ondition. Gray lines show the mean *±* SEM across these condition-level averages.

### Internal avalanche recruitment structure

#### Activation and suppression magnitudes

For each session and simulation condition, each neuron *i* had a positive, negative, or zero Avalanche Index, AI_*i*_ (for simplicity, hereafter we drop the superscript *X* indicating the simulation condition). A positive value of AI_*i*_ indicates activation of neuron *i* during the avalanche, whereas a negative value indicates suppression. For directly stimulated OV neurons, negative AI reflects gain suppression; for non-stimulated neurons, it reflects activity su ppression. To quantify the strengths of activation and suppression separately, we defined the corresponding nonnegative magnitudes:

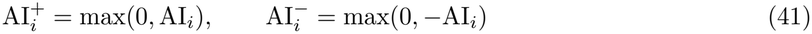

Thus, 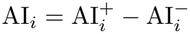, and for each neuron at most one of 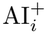 and 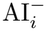 is nonzero.

#### Recruitment roles within tuning groups

For each tuning group *g* and session–offer condition, we calculated the fractions of activated and suppressed neurons as 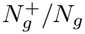 and 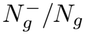, respectively, where *N*_*g*_ is the total number of neurons in the group and 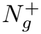 and 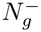 are the numbers with positive and negative AI. For groups containing at least two neurons, we quantified the fraction of neuron pairs with opposite recruitment signs as

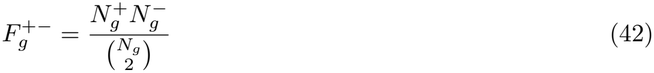

Neurons with AI = 0 contributed to *N*_*g*_ but to neither 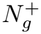 nor 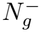. Means were computed across session–offer conditions. The 95% confidence intervals for 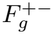 were obtained from 10,000 session-cluster bootstrap resamples, sampling sessions with replacement separately within each monkey and retaining the two offer simulations from each sampled session together.

#### Synchrony Indices

We quantified the magnitude of sign-specific pairwise recruitment within avalanches, distinguishing co-activation, co-suppression, and opposite-sign recruitment. For every unordered pair of distinct neurons (*i, j*), we defined three pairwise Synchrony Indices:

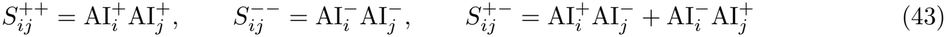

^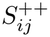^ quantifies co-activation of the two neurons, 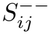 quantifies co-suppression, and 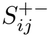 quantifies opposite-sign recruitment, corresponding to activation of one neuron and suppression of the other. Thus, “synchrony” here refers to joint recruitment within the same avalanche.

The analyses in Figs. 6 and 7 used simulations under both offers, A and B. For Fig. 6B–D, we first averaged the Synchrony Indices over neuron pairs belonging to each cell-group pair within each session– offer condition. We then plotted the mean *±* SEM of these condition-level averages across conditions. For the spatial analyses in Fig. 7, we considered only neuron pairs recorded on the same probe and defined their vertical separation as

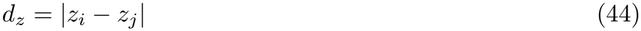

For each session–offer condition, Synchrony Indices were first computed for all same-probe neuron pairs within that condition. The resulting pair observations were then pooled across session–offer conditions to define 10 global equal-count (quantile) bins of *d*_*z*_; no neuron pairs were formed across conditions. For Fig. 7B, all pair observations falling in the same global distance bin were pooled and their Synchrony Indices were averaged. For Fig. 7CD, the same pair observations were instead grouped by recording site, and Synchrony Indices were averaged within each site and distance bin. The resulting bin-wise means were used to quantify the dependence on *d*_*z*_ across distance bins.

#### Directional Synchrony Indices and directional Bias

For the directional analysis of opposite-sign recruitment (Fig. 7EF), we further decomposed *S*^+*−*^ according to the relative *z* positions of the activated and suppressed neurons. For each same-probe pair with one neuron activated and the other suppressed, we defined the signed offset as the *z* coordinate of the activated neuron minus that of the suppressed neuron (signed *d*_*z*_ = *z*_active_ *− z*_suppressed_). Positive offsets were classified as activation superficial to suppression (sup); negative offsets as activation deep to suppression (deep); and offsets within numerical tolerance of zero as zero-offset. We partitioned the pairwise contributions to the Synchrony Index 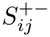 into three Directional Synchrony Indices, 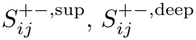, and 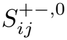, as follows. For an opposite-sign neuron pair, the full 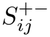 was assigned to the direction *d ∈ {*sup, deep, 0*}* determined by their signed *d*_*z*_, and zero was assigned to the other directions. If both neurons were activated, both were suppressed, or either neuron had AI = 0, the pair contributed zero to all three directions.

By construction,

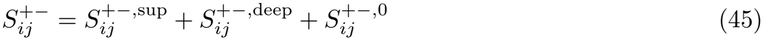

Each 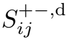, with *d ∈ {*sup, deep, 0*}*, was computed separately for each same-probe neuron pair within each session–offer condition. Same-probe pairs were assigned to the same 10 global equal-count *d*_*z*_ bins used in Fig. 7B–D, with *d*_*z*_ = |*z*_active_ *− z*_suppressed_|. Within each bin *b*, each Directional Synchrony Index was averaged across neuron pairs and session–offer conditions to obtain 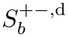. In Fig. 7E, 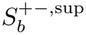 was plotted at positive *d*_*z*_ and 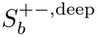 at the corresponding negative *d*_*z*_, so that the two orientations could be compared on a signed axis. We defined the directional Bias in bin *b* as

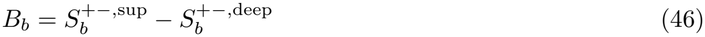

*B*_*b*_ *>* 0 indicates stronger opposite-sign recruitment when activation lies superficial to suppression (inset in Fig. 7E). To assess whether this directional preference was reproducible across recording sites (Fig. 7F), we used the largest-distance bin (1.5–2.325 mm), which also contained the maximum pooled *S*^+*−*^. For each recording site *p*, the probe-level Bias was defined as

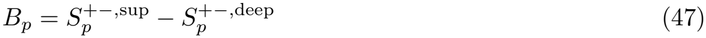

where 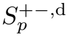 is the mean of the condition-specific 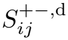 values across all neuron-pair observations from recording site *p* in the largest-distance bin.

### Statistical analysis of spatial avalanche recruitment

#### Session-bootstrap confidence intervals

We generated 10,000 bootstrap resamples by sampling recording sessions with replacement separately within each monkey, retaining each monkey’s original number of sessions. Each sampled session contributed both offer simulations, all recorded probes, and all neuron-pair observations together. Resampling whole sessions preserved dependencies among pairwise recruitment measures and between the two offer simulations. For each resample, we recomputed pooled bin means, retaining the original bin assignments. We then recalculated the Spearman correlations of mean *S*^++^ and *S*^+*−*^ with *d*_*z*_ and the fold enrichment of mean *S*^*−−*^ in the nearest-distance bin relative to all remaining pairs. The confidence intervals quantify session-resampling uncertainty conditional on the recorded monkeys and fixed distance bins; they are distinct from tests against the spatial null.

#### Recording-site scores and distance coverage

A recording site was defined by monkey identity and lateral probe coordinates (*x, y*), with pairwise recruitment measures pooled across sessions and both offers at that site. A site was included in the probe-level analyses if it contributed at least one neuron pair to at least five of the global distance bins, including the nearest and largest-distance bins. These geometry-based criteria retained 63 of 65 sites and were held fixed in every permutation (see below). Of the two excluded sites, one populated only one distance bin, insufficient to compute the correlations in Fig. 7CD; the other had no pairs in the largest-distance bin, required for the Bias analysis in Fig. 7F. At each retained site *p*, we calculated 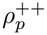 and 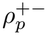 as Spearman correlations between the mean values of *S*^++^ or *S*^+*−*^ in the populated bins and the median *d*_*z*_ values of those bins in the pooled dataset. Constant profiles were assigned a score of zero by convention, indicating no distance trend, and retained in the analyses. Directional Bias *B*_*p*_ was calculated as defined above. The three component statistics were the across-site medians of 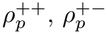, and *B*_*p*_.

#### Spatial null and component tests

We generated *M* = 10,000 spatial permutations. Within each recording session and probe, neurons’ Avalanche Indices were randomly reassigned to neuronal positions, applying the same shuffle to both offers (i.e., each neuron’s pair of responses 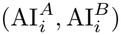 was moved together). This preserved the joint distribution of responses to the two offers and the sampling geometry within each session and probe, while disrupting the association between neuronal responses and position. After each shuffle, we recomputed the Synchrony Indices, directional assignments, site scores, and their across-site medians. The same permutations supplied the null distributions for all component and joint tests. For each component (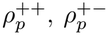 and *B*_*p*_), we compared the observed across-site median with its spatial-permutation distribution. Two-sided Monte Carlo *p*-values were computed as twice the smaller plus-one-corrected tail probability (capped at 1), with Holm correction across the three component tests.

#### Joint pattern enrichment

The joint enrichment test considered all 8 combinations of strictly positive or negative values of (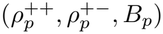, *B*_*p*_), assessing arrangements with a defined direction for all three relationships. For pattern *k*, let 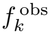 be its observed frequency among the 63 recording sites meeting the distance-coverage criteria, and *f*_*k,r*_ its frequency in permutation *r*. Sites with at least one exactly zero component are rare and belong to none of the tested patterns but remain in the denominator for both observed and shuffled frequencies. Each pattern’s expected frequency under the spatial null, 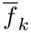, is its mean frequency across permutations. Each pattern observed enrichment is 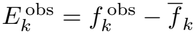 and its enrichment in permutation *r* is 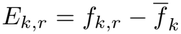. We used the largest enrichment across patterns as the joint test statistic:

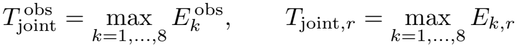

We compared the observed maximum enrichment with the distribution of maximum enrichments across spatial permutations using a one-sided Monte Carlo permutation test with the plus-one correction. Taking the maximum in both the data and each permutation accounted for selection among all 8 patterns, allowing different patterns to attain the maximum in different permutations. The complete motif is the pattern 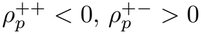, and *B*_*p*_ *>* 0. Its fold enrichment is 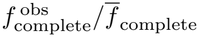. With no permutation maximum reaching the observed value, the plus-one-corrected *p*-value was 1*/*10,001, reported as 1.0*×*10^*−*4^.

#### Observed patterns and lattice schematic

For Fig. 7H, we tabulated all distinct observed patterns, including those with exactly zero components. These comprise the 8 tested patterns defined by all combinations of 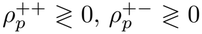, and *B*_*p*_ ≷ 0, with each sign chosen independently, and the following 3 zero-containing patterns:

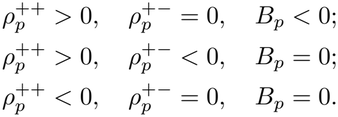

A site’s number of relationships shared with the complete motif is the number of satisfied inequalities 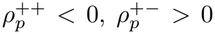, and *B*_*p*_ *>* 0; zero or opposite signs do not satisfy the corresponding relationship. Pattern frequencies use all 63 eligible sites and are grouped by the number of shared relationships, then ordered by increasing frequency in Fig. 7H, top right. The lattice schematic approximates the observed proportions sharing zero, one, two, or three relationships; site positions and absolute motif depths are illustrative.

### Tuning-corrected Synchrony Indices for spatial analysis

To test whether the spatial organization of avalanche recruitment could be explained by differences in tuning composition across distances, we constructed tuning-corrected versions of *S*^++^ and *S*^+*−*^. Each neuron pair (*i, j*) was assigned a tuning-pair label *p*(*i, j*) according to the dominant-tuning groups of its two neurons, with *p*(*i, j*) = *p*(*j, i*). The correction was performed separately for each Synchrony Index *m ∈ {*++, +*−}* and each session–offer condition *q*.

For each condition, we computed the mean Synchrony Index across all same-probe neuron pairs, pooling across *d*_*z*_,

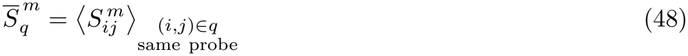

and, for each tuning-pair type *p*, the corresponding mean across same-probe pairs of that type,

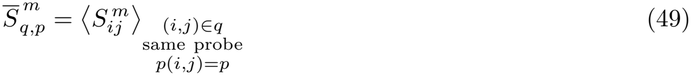

The tuning-corrected Synchrony Index was defined as

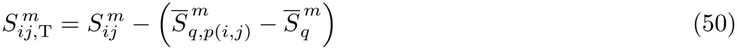

Thus, within each session-offer condition, the correction sets the mean of every tuning-pair type to the condition-wide mean across same-probe pairs. It therefore removes differences in the mean value of each Synchrony Index across tuning-pair types while preserving the condition-wide mean and pair-to-pair variation within each tuning-pair type.

The tuning-corrected spatial analyses used the same fixed 10 global equal-count *d*z bins as the corresponding uncorrected analyses. For the pooled profiles in Fig. S2AB, 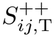 and 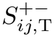 were averaged across all pair observations within each distance bin, and the dependence of the resulting bin means on *d*z was quantified by Spearman correlation. Confidence intervals were obtained with the same 10,000-sample session-cluster bootstrap used for the uncorrected profiles, resampling sessions separately within each animal and retaining both offer simulations from each sampled session together.

For the recording-site analyses in Fig. S2CD, corrected pair observations were averaged separately within each recording site and distance bin, and site-level Spearman correlations were computed from the resulting bin-wise means. The same recording-site eligibility criteria and fixed distance bins used for the uncorrected analysis were retained.

For the directional analysis in Fig. S2E, each corrected 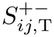 value was assigned to the active-superficial, active-deep, or zero-offset class according to the signs of the original neuron AIs and the relative *z* positions of the activated and suppressed neurons. The corrected probe-level directional Bias was then computed from the largest-distance bin using the same definition as for the uncorrected analysis.

Statistical significance was assessed with the same 10,000 within-session/probe spatial permutations used for the uncorrected analysis. For each permutation, the paired offer responses of each neuron were reassigned within the same session and probe, after which the Synchrony Indices, tuning correction, directional assignments, and recording-site statistics were recomputed. Component tests were two-sided and Holm-adjusted across the three corrected motif components.

For Fig. S2F, the joint recurrence analysis was likewise repeated using the corrected site-level signs of *ρ*^++^, *ρ*^+*−*^, and directional Bias. Enrichment of the 8 nonzero-sign patterns was assessed with the same maximum-excess permutation statistic used for the uncorrected analysis.

### Coarse-grained mean-field reductions of the inferred networks

To determine which features of the value-to-choice transformation are captured by coarse functional architecture, we constructed four reduced models from each independently inferred neuron-resolved network. In all four models, the instantaneous recurrent input generated by individual neuronal activity configurations, ∑_*j≠i*_ *J*_*ij*_*σ*_*j*_, was replaced by a self-consistent group-level mean recurrent input, ∑_*v*_ *K*_*uv*_*m*_*v*_. The reductions then varied along two orthogonal dimensions: which group-level recurrent interactions were retained and whether neuron-to-neuron heterogeneity in the intrinsic biases *h*_*i*_ was preserved. All reduced models were constructed separately for each recording session from that session’s inferred parameters; parameters were never pooled across sessions or fitted to reproduce the responses of the full network. We refer to the four reduced models as the neuron-biased functional-group mean-field model (neuron-biased FG-MF), the neuron-biased two-channel mean-field model (neuron-biased 2C-MF), the group-biased functional-group mean-field model (group-biased FG-MF), and the group-biased two-channel mean-field model (group-biased 2C-MF). The neuron-biased models retain the inferred neuron-specific intrinsic biases *h*_*i*_, whereas the group-biased models replace them by their mean *h*_*u*_ within each functional group. Independently, the FG-MF models retain all mean interactions between functional groups, whereas the 2C-MF models retain only the two offer-aligned OV–CJ interaction channels defined below. Thus, the four models form a 2*×*2 family of reductions in which interaction architecture and intrinsic-bias heterogeneity are varied independently. This design allows us to test separately the contribution of each to the value-to-choice transformation.

#### Functional-group coarse-graining of the inferred interactions

The 9 functional groups were defined by dominant neuronal tuning, following a previously described procedure [55]: OVA+, OVA*−*, OVB+, OVB*−*, CJA, CJB, CV+, CV*−*, and UC. Let *u* and *v* index these groups and let *N*_*u*_ denote the number of neurons in group *u*. For each session, we defined the mean inferred coupling between groups *u* and *v* as

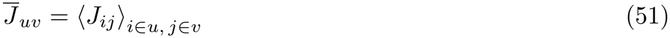

where, for *u* = *v*, the average was taken over distinct neuron pairs, *i≠j*.

For a neuron *i ∈ u*, replacing the instantaneous activity of each neuron *j ∈ v* by the mean activity *m*_*v*_ of its functional group and replacing the individual coupling *J*_*ij*_ by 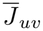 gives the recurrent input

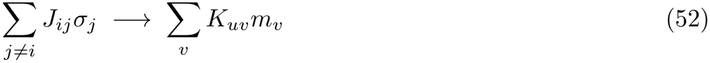

Where

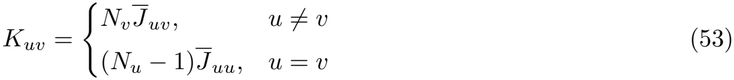

The functional-group mean-field interaction architecture retains the complete matrix *K* and therefore all mean recurrent pathways among the 9 functional groups.

#### Two-channel interaction architecture

The two-channel reduction retains only the mean OV–CJ interactions aligned with the two offer-to-choice pathways. We defined the set of retained undirected functional-group links as

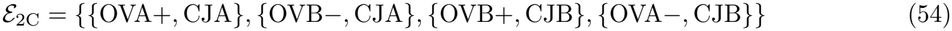

The two-channel mean interaction matrix was then (54)

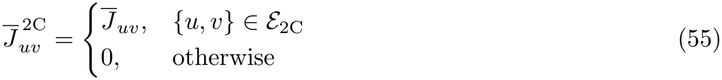

Because the underlying inferred Ising couplings are symmetric, both directions of each retained OV–CJ link are preserved. The corresponding recurrent-input matrix *K*^2C^ was obtained from Eq. (53) after replacing 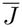 by 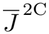. Thus, the 2C-MF models differ from the corresponding FG-MF models only in which mean functional-group interactions are retained.

#### Neuron-specific and group-mean intrinsic biases

We next varied independently whether within-group heterogeneity in the inferred intrinsic biases was retained. For each functional group *u*, we defined

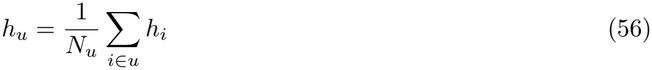

No new intrinsic-bias parameters were fitted. Instead, the intrinsic bias used for neuron *i ∈ u* was

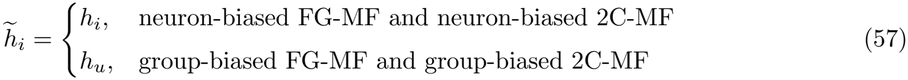

For every session and condition, the externally imposed inputs were identical to those used for the corresponding full-network simulation. In particular, we used the same baseline input, offer-specific inputs, silencing inputs for the single-offer simulations, and input strengths. Thus, differences between the full and reduced models cannot be attributed to differences in the external stimulation protocol.

#### Mean-field activities

For any reduced model *M*, let *K*(M) denote the corresponding interaction matrix, either the complete functional-group matrix *K* or the two-channel matrix *K*2*c*, and let 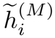 denote either *hi* or *hu*. For neuron *i* ∈ *u*, the mean-field effective input is

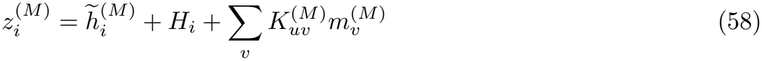

and its mean activity is

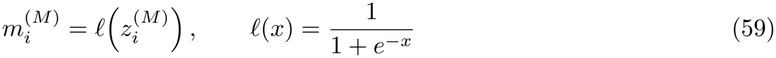

where *l*(*x*) is the logistic activation function inherited from the exact conditional activation probability of a binary neuron in the Ising model, with the instantaneous recurrent input replaced here by its mean-field approximation. The self-consistent mean activity of functional group *u* is therefore

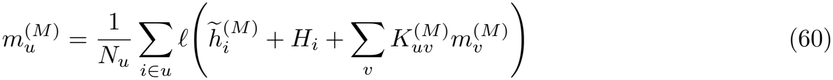

This yields 9 coupled self-consistency equations for the group-mean activities in each session and input condition. In the group-biased models, all neurons within the same functional group have the same intrinsic bias and receive the same external and mean recurrent inputs. In those models, the group-level self-consistency equations therefore simplify to

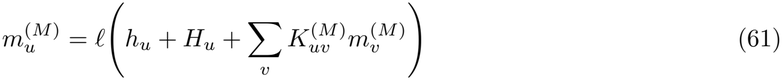

Accordingly, neurons within a given group have identical mean-field activity and susceptibility in the group-biased models, whereas neuron-biased FG-MF and 2C-MF retain neuron-specific activity and susceptibility through the individual *h*_*i*_, despite sharing the same group-level recurrent input.

The mean-field equations were solved separately for every session, model, and input condition by fixed-point iteration. For the first input value, the group-mean activities were initialized by neglecting the recurrent interaction term,

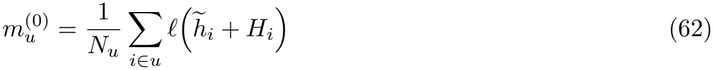

Starting from this initial condition, we iterated

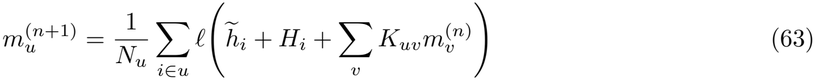

Iteration continued until

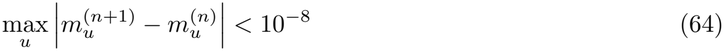

For successive nearby input values, the converged solution at the preceding input value was used as the initial condition.

### Susceptibilities and avalanche measures in the reduced models

We computed susceptibilities analytically from the mean-field equations with respect to the same input variables *X ∈ {A, B*, total*}* used for the full-network simulations. Because the externally imposed input is uniform within each functional group, the input-loading coefficient 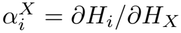, previously defined, has the same value for all neurons *i* in a given group *u*; we denote this common value by 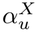. For each neuron, we defined the local mean-field gain as

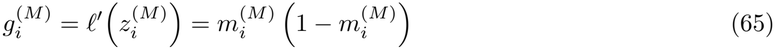

and the mean gain of functional group *u* as

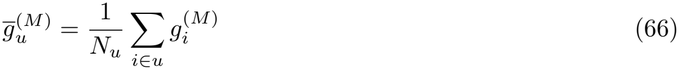

Differentiating the neuron-level mean-field equations with respect to *H*_*X*_ , the susceptibility of neuron *i ∈ u* is

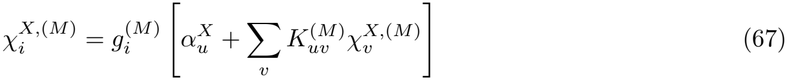

Averaging over neurons within each functional group gives

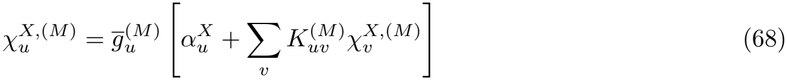

Thus, in vector form,

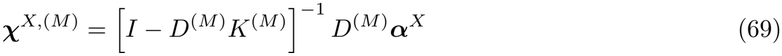

Where

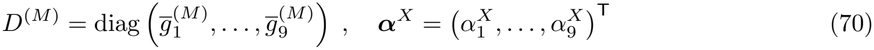

The group-level susceptibilities in Eqs. (68)–(69) were used as intermediate quantities to compute the neuron-specific susceptibilities in Eq. (67). Specifically, after solving the coupled group-level susceptibility equations, we obtained 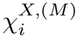 for each neuron using its neuron-specific gain 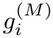. In the group-biased models, these neuron-specific susceptibilities coincide with the corresponding group-level susceptibilities, 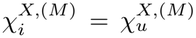 for all *i* ∈ *u*. All subsequent reduced-model analyses were based on these neuron-specific susceptibility profiles. For each neuron, the signed peak susceptibility and Avalanche Index were computed exactly as defined above for the full network, including the same subtraction of the maximal direct-input contribution 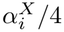 for neurons receiving a variable positive offer input. The same definitions of single-offer and competing-offer Avalanche Asymmetry were subsequently applied to the reduced-model Avalanche Indices.

#### Network mediation in the reduced models

Coarse-graining does not remove the recurrent, network-mediated nature of the response. In all reduced models, susceptibility is determined self-consistently by the retained recurrent group-level interactions, Eqs. (67), (68), and (69). These equations can be interpreted as combining the direct response to the input with its propagation through direct and indirect recurrent pathways, including recurrent loops (Supplementary Information). Even in the 2C-MF, the response remains recurrent, with OV and CJ populations influencing one another bidirectionally through the self-consistent mean-field equations.

### Aggregate value-to-choice transformation in the reduced models

For the analyses below, let *M* denote either the full inferred network or one of the four reduced models, let *s* index sessions, and let *κ* denote the input condition (*κ* = *A* or *B* for single offers and *κ* = (total, *r*) for competing offers, with *r* = *H*B/*H*A). We denote by 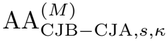 the Avalanche Asymmetry expressed in the fixed CJB ™ CJA choice coordinate defined in Eq. (39). For Fig. 5A, the mean and SEM of 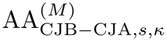 were computed across sessions separately for each of the 11 input conditions: the 2 single-offer conditions and the 9 competing-offer conditions

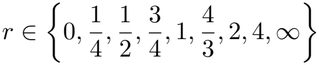

The systematic dependence of mean 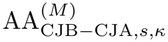 on input bias *r* = *H*_*B*_*/H*_*A*_ quantifies the aggregate value-to-choice transformation. To summarize its overall magnitude across input conditions, we generalized the offer-aligned Avalanche Asymmetry for single offers to competing-offer conditions by reversing the sign of AA_CJB*−*CJA_ for A-favoring conditions and leaving it unchanged for B-favoring conditions. The equal-offer condition *r* = 1 was excluded because neither option was favored. We refer to the resulting quantity as favored-option AA. For each model *M* , the overall magnitude of the aggregate value-to-choice transformation was defined as the mean favored-option AA across sessions and the 10 conditions with a favored option,

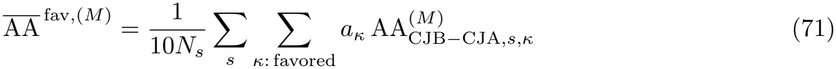

where the sum over *κ* includes the two single-offer conditions and the eight competing-offer conditions with *r≠*1. The sign *a*_*κ*_ = *−*1 for 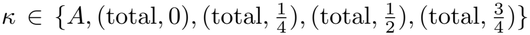 and *a*_*κ*_ = +1 for 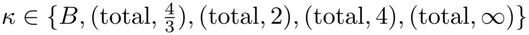, and *N*_*s*_ is the number of included sessions. For Fig. 5A, *N*_*s*_ = 60 after exclusion of the two sessions with non-converged full-network Monte Carlo simulations (as previously described) and one additional session containing a numerically converged but outlier response in the neuron-biased FG-MF. Restoring the latter session did not alter the conclusions.

### Network- and condition-specific comparison between reduced and full models

Preservation of the aggregate choice bias does not imply that a reduced model reproduces the choice bias of each inferred network under each offer condition. Because each session defines a distinct inferred network, we performed this comparison at the session–condition level.

For each of the four reduced models, we compared 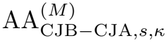 with the corresponding 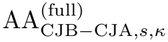 across all sessions and offer conditions, including the equal-offer condition *r* = 1.

We quantified three complementary aspects of this reduced–full model comparison. First, Spearman’s rank correlation,

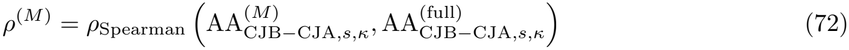

measures whether the reduced and full models preserve a common monotonic ordering of Avalanche Asymmetry across sessions and offer conditions along the A-to-B recruitment axis. This measure does not require the magnitude of Avalanche Asymmetry to be reproduced.

Second, we quantified absolute predictive agreement using

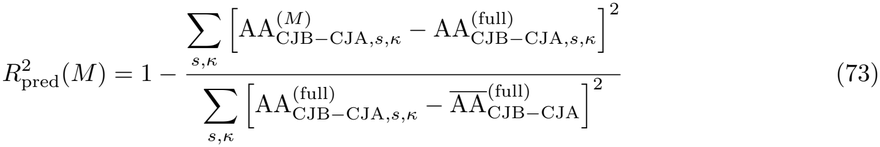

Here, 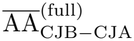 denotes the full-network Avalanche Asymmetry averaged over the same sessions and offer conditions. 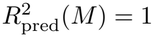indicates exact agreement between the reduced and full-network Avalanche Asymmetry, whereas 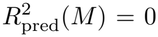 indicates that the reduced model explains none of the variation in the full-network Avalanche Asymmetry beyond simply predicting its overall mean across sessions and offer conditions. Negative values indicate performance worse than this mean-only prediction.

A reduced model can exhibit a strong Spearman correlation with the full network while having low 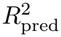: the former quantifies preservation of the rank ordering of choice biases across sessions and offer conditions, whereas the latter quantifies how accurately their magnitudes are reproduced.

Third, we quantified the typical absolute response error as

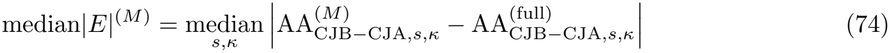

This measure is expressed directly in Avalanche Asymmetry units and complements 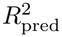.

For Fig. 5B, the outlier response in the neuron-biased FG-MF was shown but excluded from the main statistics; restoring this observation made 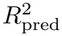 negative for that model but did not substantially alter the other statistics or the conclusions.

### Neuron-resolved recruitment in the neuron-biased FG-MF

To determine whether the systematic neuron-resolved recruitment structure observed in the full networks can already emerge from coarse functional-group interactions, we focused on the neuron-biased FG-MF. We used this model for two reasons. First, it retains neuron-specific biases and can therefore generate heterogeneous recruitment among neurons within the same functional group, allowing it in principle to reproduce the within-group activation–suppression subdivision and the associated neuron-resolved spatial organization. Second, among the four reduced models, the neuron-biased FG-MF is also the least compressed description and, after exclusion of the outlier, shows the lowest median |*E*| and highest 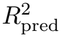. Accordingly, to assess neuron-resolved recruitment in the neuron-biased FG-MF (Fig. S3), we applied the same recruitment-role analysis as in Fig. 6A and the same probe-level spatial-lattice analyses and spatial-permutation tests as in Fig. 7CDFG. The session containing the outlier response was excluded; including it did not alter the conclusions.

## Author Contributions

GT designed the study. YK and CPS conducted the experiment and performed spike sorting and cell classification; GT and TGC curated an automated software package to run the network-inference algorithm on the data; KZ performed all Monte Carlo simulations under single and competing offers, the ablation analyses, and quantification of the results; GT performed the analyses of neuron-resolved recruitment, synchrony, and lattice structure. GT and KZ derived and analyzed the mean-field model reductions. All the authors discussed and evaluated the results. KZ and GT wrote the initial draft; all the authors edited the manuscript. CPS directed the experimental component; GT directed the theoretical and computational components of the project.

## Acknowledgments

This work was supported by the Sloan Foundation (Research Fellowship Award No. FG-2024-22163 to GT), the McDonnell Center for Systems Neuroscience (Small Grant GF0013150 to GT), the NIH (grants R01-MH104494 and R01-DA032758 to CPS), the Japan Student Services Organization (predoctoral fellowship to YK), Washington University (BioSURF Fellowship Award for undergraduate research to TGC), and the Abdus Salam International Centre for Theoretical Physics (Rita Levi-Montalcini Post-doctoral Fellowship to KZ).

