## Supplementary Information for "Inferred interactions transform values into lattice-patterned choice attractors in orbitofrontal cortex"

#### Monte Carlo convergence diagnostics and validation

##### Diagnostic procedure and exclusion criterion

Each Monte Carlo simulation, corresponding to a fixed inferred network and a fixed external input, consisted of a sequence of pre-run windows followed by a separate production window. The pre-run windows were used to assess whether Monte Carlo estimates had stabilized, whereas the production window provided the samples used to compute the final observables and served as an additional convergence check. We assessed convergence independently for neuronal mean activity, $\langle \sigma_i \rangle$, and susceptibility, $\chi_i$, as defined in the Methods. Because convergence was evaluated separately for each simulation condition, we suppress the corresponding condition label $X \in \{A, B, \text{total}\}$ throughout this section. For compactness, we use $q_i$ to denote either observable,

$$q_i \in \{\langle \sigma_i \rangle, \chi_i\}$$

Neurons deterministically silenced by the imposed negative input were excluded from the convergence calculations because their state remained fixed at $\sigma_i = 0$ throughout the simulation. Their mean activity and susceptibility therefore contained no information about equilibration of the Monte Carlo trajectory.

**Pre-run windows and Monte Carlo standard errors.** For each simulation, we used consecutive, non-overlapping pre-run windows of length

$$W_t \in \{2^{14}, 2^{15}, \ldots, 2^{20}\} \text{ MC sweeps}$$

All simulations were run through the full pre-run schedule; convergence was therefore assessed diagnostically rather than used as an early-stopping criterion.

*Co-senior authors

Within each window, one network state was collected every $q_{\text{save}} = 5$ MC sweeps. Because successive sampled states can retain residual Monte Carlo correlations, standard errors were estimated by blocking rather than by treating all sampled states as independent. The samples in each window were divided into consecutive, non-overlapping blocks of $L = 100$ states, corresponding to 500 MC sweeps per block. The number of complete blocks in window $t$ was $B_t = \lfloor \lfloor W_t/q_{\text{save}} \rfloor / L \rfloor$. Within each block, mean activity and susceptibility were recomputed using the same estimators as for the full simulation window. In particular, for susceptibility, the covariance-based estimator was recomputed using the means estimated within that block rather than means from the full window. The block estimates were used only to quantify sampling uncertainty; the point estimate $\hat{q}_i(t)$ entering the convergence analysis was always the estimate obtained from the entire pre-run window.
For either observable, the window-level Monte Carlo standard error was estimated as

$$SE_i(t) = \frac{s_{i,\text{block}}(t)}{\sqrt{B_t}}$$

where $s_{i,\text{block}}(t)$ is the sample standard deviation of the corresponding blockwise estimates. This procedure accounts more conservatively for residual autocorrelation than a standard error obtained by dividing the single-sample standard deviation by the square root of the total number of sampled states.

**Between-window discrepancy.** For each non-silenced neuron $i$, we compared the estimates from the final two pre-run windows. We defined

$$\Delta_i = |\hat{q}_i(t) - \hat{q}_i(t-1)| \tag{S1}$$

$$A_i = \max\left(|\hat{q}_i(t)|, |\hat{q}_i(t-1)|\right) \tag{S2}$$

$$SE_{i,\Delta} = \sqrt{SE_i(t)^2 + SE_i(t-1)^2} \tag{S3}$$

Here, $\Delta_i$ is the observed change between the two estimates, $A_i$ provides a neuron-specific scale for the observable, and $SE_{i,\Delta}$ quantifies the Monte Carlo sampling uncertainty in the between-window difference. We allowed a small relative tolerance, $\epsilon = 0.02$, and defined the normalized discrepancy

$$R_i = \frac{\Delta_i - \epsilon A_i}{SE_{i,\Delta}}$$

Thus, $R_i$ quantifies the between-window discrepancy beyond that expected from Monte Carlo sampling variability, after allowing for a 2% relative tolerance. Larger $R_i$ therefore indicates stronger evidence that the apparent change reflects residual lack of equilibration rather than ordinary sampling noise. A neuron was flagged when

$$R_i > 2$$

or, equivalently,

$$\Delta_i > \epsilon A_i + 2SE_{i,\Delta}$$

The $R_i > 2$ threshold was used as a sensitive neuron-level diagnostic rather than as a requirement that every neuron pass. Under a stationary Gaussian sampling approximation, and in the limiting case in which $\epsilon A_i$ is negligible – which is also the regime in which the diagnostic is most sensitive to discrepancies – the nominal two-sided probability of obtaining $R_i > 2$ from finite-sampling fluctuations alone is

$$2[1 - \Phi(2)] \simeq 0.0455$$

where $\Phi$ is the cumulative distribution function of the standard normal distribution. Thus, approximately 4.55% of neuron-level estimates can exceed this threshold by chance even after equilibration. For non-negligible $\epsilon A_i$, the exceedance probability is smaller.

**Extreme discrepancies.** We next asked whether a discrepancy was unusually large after accounting for the number of neurons examined in a simulation. For a simulation containing $N$ non-silenced neurons, we defined the Bonferroni family-wise threshold

$$R_{\text{extreme}} = \Phi^{-1}\left(1 - \frac{0.05}{2N}\right)$$

Under the stationary Gaussian reference, this threshold limits the nominal probability that at least one of the $N$ neurons exceeds the threshold by chance to at most 5%. We therefore classified

$$R_i \geq R_{\text{extreme}}$$

as an extreme discrepancy. The two thresholds distinguish ordinary sampling-tail events from unusually large discrepancies: $R_i > 2$ provides a sensitive neuron-level diagnostic, whereas $R_i \geq R_{\text{extreme}}$ identifies discrepancies warranting further examination.

**Persistent extreme discrepancies.** After completion of the full pre-run, the same Markov chain was continued into a new, non-overlapping production window. The production window contained $N_{\text{prod}} =$ 20,000 sampled states, with one state recorded every 5 MC sweeps, corresponding to 100,000 additional MC sweeps. The final pre-run window was not reused as part of the production sample. The production mean-activity and susceptibility estimates were those used in the subsequent analyses.
Production-window standard errors were obtained using the same block procedure described above. With blocks of 100 sampled states, the production window contained $B_{\text{prod}} = 200$ blocks.
For every neuron showing an extreme discrepancy in the final pre-run comparison, we compared the final pre-run estimate with the production estimate using

$$R_i^{\text{prod}} = \frac{|\hat{q}_i^{\text{prod}} - \hat{q}_i(t)| - \epsilon \max\left(|\hat{q}_i^{\text{prod}}|, |\hat{q}_i(t)|\right)}{\sqrt{(SE_i^{\text{prod}})^2 + SE_i(t)^2}}$$

An extreme discrepancy that did not recur in this comparison was consistent with an isolated finite-sampling fluctuation. We defined a persistent extreme discrepancy as one satisfying

$$R_i \geq R_{\text{extreme}} \qquad \text{and} \qquad R_i^{\text{prod}} \geq R_{\text{extreme}}$$

**Residual drift.** As an independent diagnostic of incomplete equilibration, we tested for systematic directional drift near the end of each simulation. For every neuron, we considered the sequence

$$\hat{q}_i(t-2),\ \hat{q}_i(t-1),\ \hat{q}_i(t),\ \hat{q}_i^{\text{prod}}$$

formed by the final three pre-run windows and the production window, and determined whether it was monotonically increasing or monotonically decreasing.
The observed fraction of monotonic trajectories was compared with that expected under a stationary independent-Gaussian null, with the unequal standard errors of the four estimates taken into account. An excess of monotonic trajectories would therefore indicate systematic residual drift rather than ordinary sampling variability.

**Condition-level convergence.** Finally, to assess convergence across the complete input sweep of each network–stimulation condition, we introduced two condition-level diagnostics quantifying the prevalence and severity of extreme discrepancies. A condition $c$ denotes one fixed network realization under one fixed stimulation protocol, simulated across all input values (an H-sweep). This definition applies to inferred networks, ablated networks, matched random controls, and independently generated random networks.
For observable $q$ and condition $c$, we defined

$$p_c^{(q)} = \frac{\#\left\{H \in \mathcal{H}_c : \begin{array}{c}\text{at least one non-silenced neuron}\\ \text{has } R_{i,c}^{(q)}(H) \geq R_{\text{extreme}}(N_c)\end{array}\right\}}{\#\mathcal{H}_c}, \qquad p_c = \max_q p_c^{(q)}$$

where $\mathcal{H}_c$ is the set of evaluated input values, $N_c$ is the number of non-silenced neurons in condition $c$, and $q$ denotes susceptibility or mean activity. Thus, $p_c$ is the maximum, across susceptibility and mean activity, of the fraction of input values $H$ with at least one extreme neuronal discrepancy.
To assess whether extreme discrepancies persisted and affected substantial fractions of neurons at any input value, we also defined the *maximum persistent-discrepancy fraction*, $F_c$. Let $n_{c,\text{pers}}^{(q)}(H)$ denote the number of non-silenced neurons exhibiting a persistent extreme discrepancy in observable $q$ at input $H$. Specifically, a neuron is counted only if its discrepancy meets or exceeds $R_{\text{extreme}}(N_c)$ in both comparisons:

between the final two pre-run windows and between the final pre-run and production windows. We computed

$$F_c^{(q)} = \max_{H \in \mathcal{H}_c} \frac{n_{c,\mathrm{pers}}^{(q)}(H)}{N_c}, \qquad F_c = \max_q F_c^{(q)}$$

Thus, $F_c$ measures the largest fraction of persistently discrepant neurons at any input value and in either observable. Instability affecting many neurons at only a few input values can produce a modest $p_c$, whereas $F_c$ captures the fraction of affected neurons.

**Operational exclusion criterion.** We computed one $F_c$ for every condition defined above and pooled these condition-level values across all single-offer, competing-offer, ablation, and random-network simulations before any convergence-based exclusions. In this dataset, the pooled distribution of $F_c$ is bimodal, with a dominant mode at 0% and a second mode near 100%. We therefore set the exclusion cutoff at $F_{\mathrm{cut}} = 50\%$, separating the two modes, and excluded all conditions with $F_c > F_{\mathrm{cut}}$. The classification is largely insensitive to the specific value of this cutoff. Changing $F_{\mathrm{cut}}$ from 10% to 60% reclassifies only 0.18% of the conditions. After applying this exclusion rule, the remaining convergence diagnostics – the frequencies of ordinary between-window discrepancies ($R_i > 2$) and Bonferroni-extreme discrepancies ($R_i \geq R_{\mathrm{extreme}}$), the persistence of extreme discrepancies in the production-window comparison, residual drift, and the condition-level distribution of $p_c$ – were used to validate the resulting classification.

#### Convergence validation results

Together, the convergence diagnostics provide strong evidence that all retained simulations had equilibrated, while clearly separating the very few excluded conditions as exceptional cases of persistent instability. Applying the $F_c$ exclusion rule identified both single-offer conditions from session D240215c as non-converged ($F_c > F_{\mathrm{cut}}$); this session was therefore excluded from all subsequent simulation analyses. In the competing-offer simulations, multiple conditions from session D240411c also exceeded $F_{\mathrm{cut}}$, and this session was excluded from that analysis. No further exclusions were required for the ablation simulations, including the randomly ablated controls, or for random controls matched to the retained inferred networks. Among the independently generated random networks in Fig. 2D, some conditions exceeded the exclusion criterion and were removed; only retained conditions contributed to that figure. The convergence diagnostics validating these classifications are reported below.

##### Single-offer simulations

Unless otherwise stated, convergence statistics were computed across the 124 session×offer conditions remaining after exclusion of session D240215c (4,216 simulations).

1. **Between-window discrepancies occur at rates compatible with finite sampling.**

   The fraction of non-silenced neurons with $R_i > 2$ is 3.78% for susceptibility and 0.52% for mean activity. Both are below the nominal Gaussian upper reference $2[1 - \Phi(2)] = 4.55\%$. Thus, the observed differences between consecutive pre-run windows are compatible with ordinary finite-sampling fluctuations rather than systematic lack of equilibration.

2. **Extreme discrepancies are rare and consistent with the stationary Gaussian null.**

   4.25% of simulations contain at least one neuron with an extreme susceptibility discrepancy, whereas 0.26% contain at least one neuron with an extreme mean-activity discrepancy. Both fractions are below the nominal 5% Bonferroni family-wise upper bound expected under the stationary Gaussian reference.

3. **Extreme discrepancies almost never persist in the separate production window.**

   Persistent extreme discrepancies account for only 0.0031% of susceptibility estimates and 0% of mean-activity estimates. Only 7 persistent susceptibility instances are observed in total. They occur in 6 session×offer conditions, while the remaining 118 conditions contain none. Thus, apparent extreme discrepancies observed in the pre-run almost always disappear when tested against the separate production window.

4. **There is no evidence of systematic residual drift.**

   The fractions of neuron-level trajectories exhibiting monotonic drift are 14.64% for susceptibility and 14.50% for mean activity. These values closely match the 14.46% expected under the stationary independent-Gaussian null after accounting for the unequal standard errors of the different windows.

5. **The exclusion criterion clearly identifies the exceptional non-converged session.**

   Among the 124 retained network–stimulation conditions, $F_c = 0$ for 95.2% of conditions and the maximum is 3.70%. In contrast, both single-offer conditions associated with D240215c have $F_c > F_{\text{cut}}$, with $F_c$ reaching 87.88%. Condition-level extreme discrepancies, quantified by $p_c$, are likewise rare among retained conditions but elevated in D240215c. At the neuron level, approximately 42–48% of susceptibility estimates in D240215c have $R_i > 2$, 33–37% have $R_i \geq R_{\text{extreme}}$, and approximately 24–30% show persistent extreme discrepancies.

##### Competing-offer simulations

After excluding D240215c, the $F_c$ criterion identifies D240411c as an additional exceptional non-converged session, with most of its conditions exceeding $F_{\text{cut}}$. Among the conditions from the retained sessions, $F_c = 0$ for 95.8% of them and the maximum is 30.88%. In contrast, $F_c$ reaches 88.79% in D240411c.
The remaining convergence diagnostics independently support the classification of D240411c as an exceptional case with substantial residual sampling instability and provide strong evidence of equilibration in the remaining competing-offer simulations. Specifically, across the retained conditions, 3.45% of susceptibility estimates have $R_i > 2$ and only 3.59% of simulations contain at least one extreme neuron, both below their nominal stationary-Gaussian reference levels of 4.55% and 5%, respectively. Only 0.15% of retained simulations contain at least one persistent extreme neuron, and the fraction of susceptibility trajectories showing monotonic drift is 14.43%, closely matching the 14.46% expected under the stationary independent-Gaussian null. Mean activity likewise shows no evidence of lack of convergence: 0.45% of estimates have $R_i > 2$, only 0.23% of retained simulations contain at least one extreme neuron, and 14.42% of trajectories show monotonic drift. Condition-level extreme discrepancies quantified by $p_c$ are likewise concentrated at low values among retained conditions.

##### Ablation simulations

After excluding session D240215c, all remaining targeted- and random-ablation conditions have $F_c < F_{\text{cut}}$ and are retained. Condition-level extreme discrepancies quantified by $p_c$ are likewise rare across the retained targeted- and random-ablation conditions. Both the overall fraction of neurons with $R_i > 2$ and the fraction of simulations containing at least one neuron with $R_i \geq R_{\text{extreme}}$ are below their corresponding nominal chance levels.

##### Random-network controls

Random-network controls were assessed using the same convergence diagnostics as the inferred networks. Each condition $c$ comprised a complete H-sweep for one random network realization under one fixed stimulation protocol. We computed $F_c$ and $p_c$ for each condition and assessed convergence separately for random controls matched to the retained inferred networks and for the independently generated random networks in Fig. 2D.
For the matched random controls, all conditions have $F_c < F_{\text{cut}}$ and are retained. Their $p_c$ distributions are likewise concentrated at low values, consistent with equilibration.
No conditions are excluded for independently generated random networks with mean $J_{ij} = 0.010$ or 0.026. At mean $J_{ij} = 0.050$, 342 of 5,500 conditions have $F_c > F_{\text{cut}}$ and are excluded. Non-convergence is concentrated in the largest networks: approximately 54% of realizations with $N = 150$ fail to converge, compared with 14% at $N = 137$ and only two realizations at $N = 125$. Only retained conditions contribute to Fig. 2D. The $p_c$ distributions and the remaining convergence diagnostics independently support this classification.

### Susceptibility Peak Alignment in inferred and random networks

Although random recurrent connectivity can also produce coordinated susceptibility responses, Alignment was significantly higher in inferred networks ($p = 0.0054$, two-sided Wilcoxon signed-rank test; Fig. S1). In each of the three monkeys, larger Alignment differences tended to occur in conditions with elevated recruitment of CJ neurons, further linking susceptibility-peak coordination to the value-to-choice transformation.

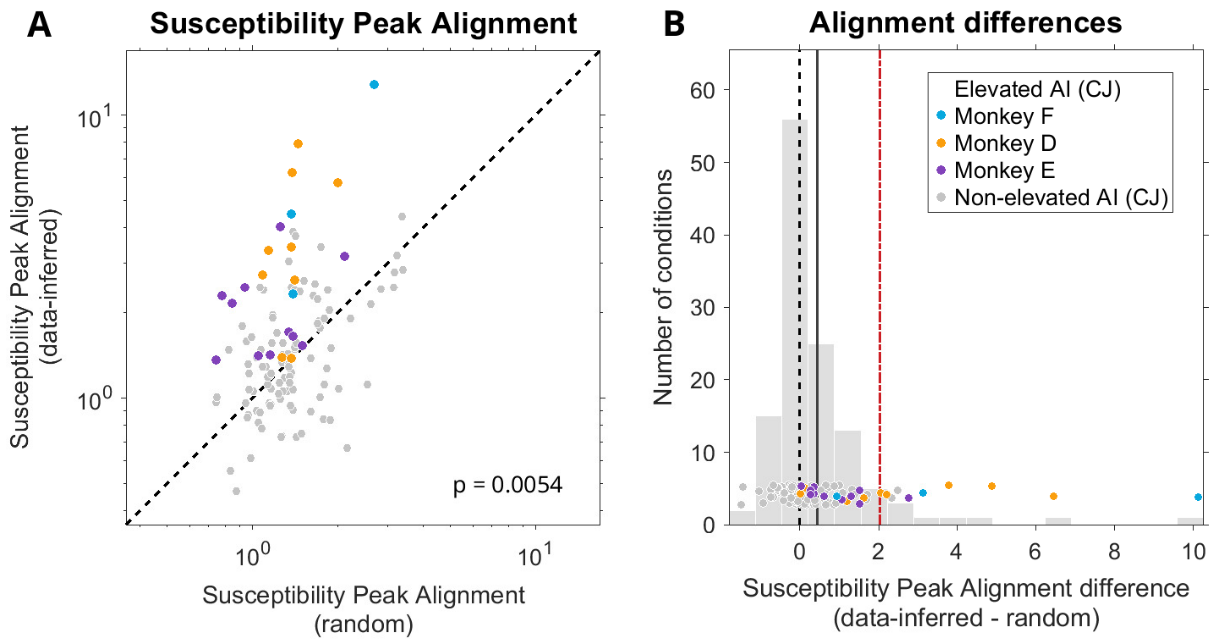


Figure S1: **Susceptibility Peak Alignment in inferred and matched random networks. (A)** Susceptibility Peak Alignment in inferred networks versus matched random networks across session–offer conditions. For each condition, random-network Alignment was defined as the inverse of the mean susceptibility-weighted peak dispersion across 10 matched random realizations. The dashed line indicates equality. Colors identify conditions with elevated CJ Avalanche Index (AI) in each monkey; gray indicates conditions with non-elevated CJ AI (as in Fig. 2C). Alignment was significantly higher in inferred than in random networks ($p = 0.0054$, two-sided Wilcoxon signed-rank test across all session–offer conditions). **(B)** Distribution of Alignment differences (data-inferred minus random) across the same conditions. The dashed vertical line indicates zero difference, the solid gray line indicates the mean difference across all conditions, and the dash-dotted red line indicates the mean difference across conditions with elevated CJ AI (colored data points). Individual conditions are shown as data points using the same color scheme as in (A). Larger positive differences tend to occur in conditions with elevated mean AI of CJ neurons, linking stronger value-to-choice recruitment to a more coordinated population response.

### Lattice organization persists after controlling for tuning composition

We repeated the spatial analyses in Fig. 7 using the tuning-corrected Synchrony Indices defined in Methods. The pooled distance profiles, recording-site recurrence analyses, directional Bias, and joint motif-enrichment analysis were recomputed using the same distance bins, eligibility criteria, bootstrap procedure, and spatial-permutation procedure. The tuning-corrected results are shown in Fig. S2 and demonstrate that the characteristic lattice relationships persist after controlling for tuning composition.

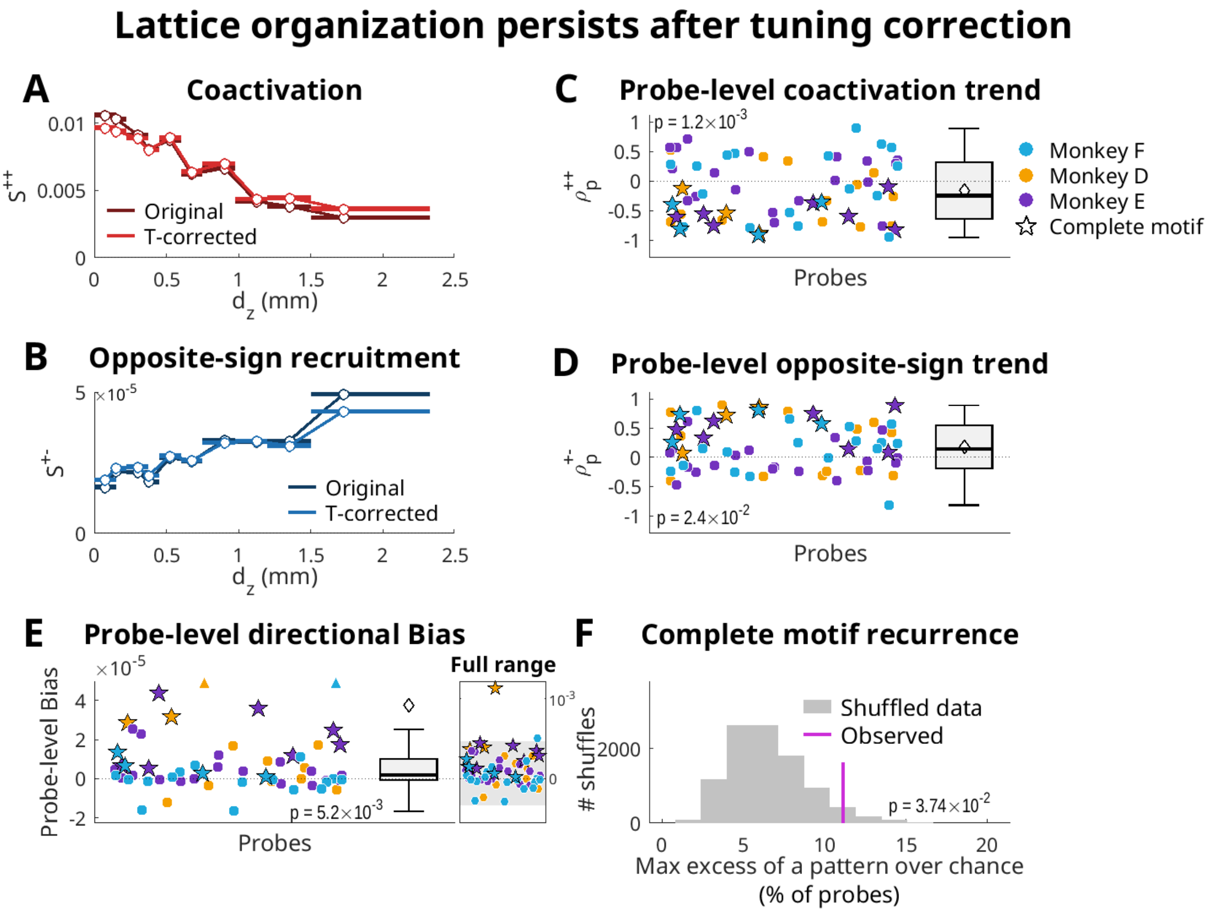


Figure S2: **Lattice organization persists after controlling for tuning composition.** (**A,B**) Pooled same-probe spatial profiles of (A) co-activation, $S^{++}$, and (B) opposite-sign recruitment, $S^{+-}$, before tuning correction (Original) and after tuning correction (T-corrected). Pair observations were assigned to the same 10 global equal-count $d_z$ bins used in Fig. 7; points show the mean Synchrony Index within each bin, and horizontal bars indicate the corresponding $d_z$ range. (**C,D**) Recording-site Spearman correlations between $d_z$ and (C) tuning-corrected $S^{++}$ or (D) tuning-corrected $S^{+-}$. (**E**) Recording-site directional Bias for tuning-corrected $S^{+-}$ in the largest-distance bin (positive values indicate stronger opposite-sign recruitment when activation is superficial to suppression). The inset shows the full range of Bias values. In (C–E), points denote individual recording sites and colors indicate monkeys; stars mark sites expressing the complete motif in the tuning-corrected scores. Box plots show the median and interquartile range, whiskers extend to the outermost data point within 1.5× the interquartile range, and diamonds indicate the mean. Reported $p$ values compare the median recording-site Spearman correlations in (C,D) and directional Bias in (E) against their respective spatial-permutation null distributions; all tests are two-sided and Holm-adjusted across the three components. (**F**) Joint recurrence of the complete motif after tuning correction. Gray bars show the spatial-permutation distribution of the maximum excess in pattern frequency relative to its shuffled expectation, evaluated across all 8 possible sign patterns. The tuning correction was recomputed for each permutation. The magenta line shows the observed maximum excess, 11.1 percentage points, corresponding to the complete motif ($p = 0.0374$).

#### Interaction coarse-graining disrupts neuron-resolved recruitment structure

To test whether the systematic neuron-resolved recruitment patterns observed in the full networks arise from coarse interactions between tuning groups, we analyzed the neuron-biased FG-MF, which retains neuron-specific biases $h_i$ but replaces neuron-specific interactions with tuning-group means. This reduction does not preserve the systematic subdivision of tuning groups into activated and suppressed neurons, and neither the three probe-level lattice components nor the complete motif differ significantly from the spatial null (Fig. S3). We performed the analysis with and without session F241105c, whose single-offer B condition produced extreme near-singular amplification in the neuron-biased FG-MF. Fig. S3 shows the results excluding this session; the conclusions were unchanged when it was included.

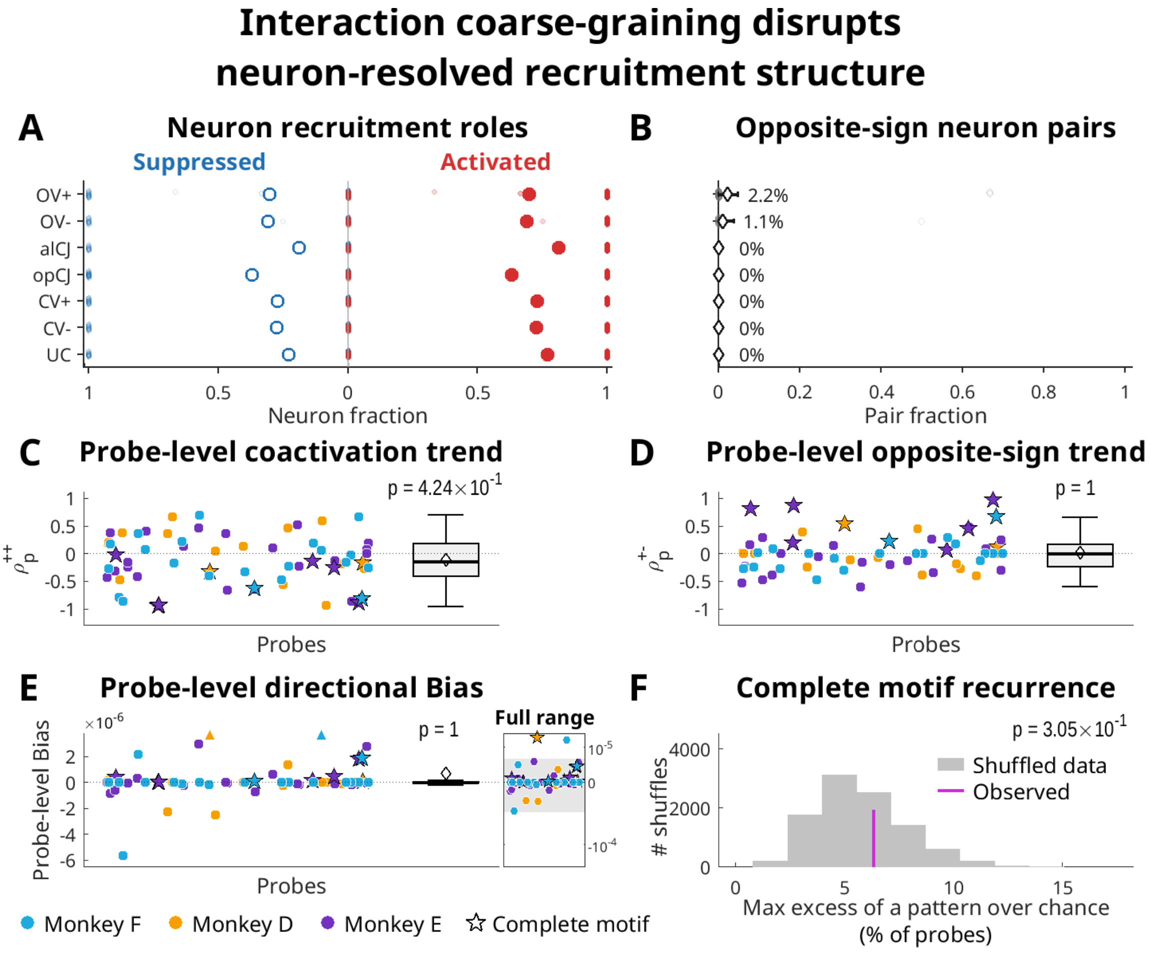


Figure S3: **Interaction coarse-graining disrupts neuron-resolved recruitment structure.** Neuron-resolved recruitment in the neuron-biased functional-group mean-field model (FG-MF), which retains the inferred neuron-specific biases $h_i$ while replacing neuron-specific interactions with tuning-group means. **(A)** Fractions of activated and suppressed neurons within each tuning group. Small points show individual session–offer conditions and large points show means, as in Fig. 6A, left. Recruitment within each group is nearly uniform across neurons in individual conditions, with fractions concentrated near 0 or 1, indicating loss of the systematic activation–suppression splitting observed in the full networks. The nonzero mean fractions largely reflect variation in recruitment sign across conditions rather than coexistence of activated and suppressed neurons within the same condition. **(B)** Fraction of opposite-sign neuron pairs within each tuning group and session–offer condition; diamonds show means across conditions and horizontal bars show 95% session-bootstrap confidence intervals, as in Fig. 6A, right. Mean fractions are near zero across all tuning groups ($\leq 2.2\%$), confirming that systematic within-group activation–suppression splitting is largely lost after interaction coarse-graining. **(C–E)** Probe-level components of the spatial lattice: distance dependence of coactivation, $\rho_p^{++}$ **(C)**; distance dependence of opposite-sign recruitment, $\rho_p^{+-}$ **(D)**; and directional Bias **(E)**. Plotting conventions and statistical tests are as in Fig. 7CDF. None of the three lattice components differs significantly from the spatial null. **(F)** Joint recurrence of the complete lattice motif. Plotting conventions and statistical test are as in Fig. 7G. The complete motif does not differ significantly from the spatial-permutation null.

#### Recurrent-path interpretation of mean-field susceptibility

Eqs. (68) and (69) show how the group-level susceptibility is generated by the externally imposed input and its recurrent propagation through the retained functional-group interactions. The input-loading vector $\boldsymbol{\alpha}^X$ specifies how the external input to each functional group changes as $H_X$ is varied. The matrix $D^{(M)}$ determines the local sensitivity of each group at the current mean-field operating point: groups whose neurons lie in a high-gain region of their input–output functions respond more strongly to a given change in effective input. The factor

$$\left[I - D^{(M)} K^{(M)}\right]^{-1}$$

then describes how these local responses are propagated through the recurrent interactions retained in model $M$. For simplicity, we suppress the model superscript $(M)$ below. At a fixed operating point, when the spectral radius satisfies $\rho(DK) < 1$ (i.e., when successive rounds of propagation through the network progressively decay), the inverse admits the convergent Neumann expansion

$$(I - DK)^{-1} = I + DK + (DK)^2 + (DK)^3 + \cdots \tag{S4}$$

Consequently,

$$\boldsymbol{\chi}^X = D\boldsymbol{\alpha}^X + DKD\boldsymbol{\alpha}^X + DKDKD\boldsymbol{\alpha}^X + \cdots \tag{S5}$$

The first term is the direct response of the externally driven groups. The second term contains responses propagated through one retained group-to-group interaction, the third contains responses propagated through two successive interactions, and higher-order terms contain progressively longer recurrent paths, including recurrent loops. Thus, the reduced models retain self-consistent propagation through direct and higher-order recurrent pathways, within a coarse-grained interaction architecture.
